# An explainable map of human gastruloid morphospace reveals gastrulation failure modes and predicts teratogens

**DOI:** 10.1101/2024.09.20.614192

**Authors:** Joseph Rufo, Chongxu Qiu, Dasol Han, Naomi Baxter, Gabrielle Daley, Maxwell Z. Wilson

## Abstract

Human gastrulation is a critical stage of development where many pregnancies fail due to poorly understood mechanisms. Using the 2D gastruloid, a stem cell model of human gastrulation, we combined high-throughput drug perturbations and mathematical modelling to create an explainable map of gastruloid morphospace. This map outlines patterning outcomes in response to diverse perturbations and identifies variations in canonical patterning and failure modes. We modeled morphogen dynamics to embed simulated gastruloids into experimentally-determined morphospace to explain how developmental parameters drive patterning. Our model predicted and validated the two greatest sources of patterning variance: cell density-based modulations in Wnt signaling and SOX2 stability. Assigning these parameters as axes of morphospace imparted interpretability. To demonstrate its utility, we predicted novel teratogens that we validated in zebrafish. Overall, we show how stem cell models of development can be used to build a comprehensive and interpretable understanding of the set of developmental outcomes.

## Introduction

Human gastrulation, which involves two symmetry breaking events and the specification of the germ layers, has been hypothesized to be a go, no-go decision for the embryo– acting as an early test of the fidelity of developmental processes that are essential for proper formation later.^1,2^ Indeed, as many as 30% of human pregnancies are predicted to fail at this critical stage,^3,4^ yet because of anatomical differences between human and other vertebrate models of gastrulation,^5^ as well as ethical constraints on experimentation with human embryos,^6^ we lack a comprehensive view of the key parameters that govern human gastrulation and how perturbations to these parameters lead to failure.

Inborn errors of gastrulation can lead to either spontaneous abortion or congenital malformations.^7,8^ In fact, there are more deaths attributed to errors of development than all pediatric cancers^9^ and the mortality rate attributed to congenital malformations is higher than deaths caused by stroke, Alzheimer’s disease, and diabetes combined.^10,11^ These trends have persisted over the past 15 years,^12^ with little progress in reducing mortality rates or advancing our ability to prevent or treat congenital malformations. This problem persists, because the causes of congenital malformations remain poorly understood, with 50-80% having no known etiology.^13,14^ Despite this public health burden, state-of-the-art methods to assess the teratogenicity of compounds have not advanced beyond exposing model organisms, such as rats, mice, and zebrafish, to potential teratogens and assessing gross changes in morphology. Notably, such models do not capture the human-specific teratogens–mice and rats are notoriously resistant to thalidomide-induced birth defects^15,16^–and therefore constitute a dire need for quantitative methods to predict, even minor harm to the developing embryo.

Unbiased characterization of state-space manifolds, in the context of development referred to as morphospaces^17^ are an emerging set of methods that show promise for understanding the various ways in which development can fail, how to predict these failures *a priori*, and, potentially, open up avenues for treatment. Three key experimental characteristics are essential to build a morphospace representation: (1) a quantitative, high-dimensional measurement of the system state, (2) a reproducible set of initial conditions, and (3) a diverse and unbiased set of perturbations. Such methods allow for a “first principles” style approach to understanding the set of possible system states and their inter-relationships and offer a framework for organizing the vast number of possible phenotypes in a manner that allows for quantitative models that can both predict the effects of perturbations on embryonic development and suggest what system parameters can be targeted to avoid such errors.

The rise of synthetic models of human development have generated a number of engineered systems with which to explore human morphospace.^18^ Stem cell-based, 3D models including post-gastrulation structured stem-cell-based embryo models (SEMs),^19^ post-implantation embryoids,^20^ peri-gastruloids,^21^ post-implantation amniotic sac embryoids (PASE),^22^ blastoids,^23^ gastruloids,^24^ and somitoids,^25^ represent distinct developmental events and differing in their ability to recapitulate anatomical structure, morphological organization, and function. However, 3D embryo models suffer from high levels of variability and low formation efficiencies (0.42% of starting aggregates possess proper structural organization and morphology),^19^ and therefore, under current protocols, would require the generation of an impractical number of constructs for high-throughput screening. On the other hand, 2D^26^ or 2.5D^27^ models that leverage lithographic micropatterned surfaces achieve near uniform formation of 10^3^-10^4^ constructs per experiment,^28^ albeit at the cost of altering the topology of the developmental tissue. Of note, the 2D gastruloid, a micropatterned disc of embryonic stem cells is an exceptional model of the human primitive streak noted for its ability to reproducibly differentiate into radially patterned germ layers and recreate all cell types of the gastrulating human embryo.^29^ The 2D gastruloid is thus a prime candidate model with which to map the morphospace of human gastrulation.

Despite the potential advantages of 2D models in studying human morphospace, capturing the full complexity of these systems requires innovative computational approaches to determine underlying relationships in the data. Linear covariance-based methods, such as principal component analysis (PCA), are sensitive to scaling and struggle with datasets where the relationships between variables are non-linear. On the other hand, more recently developed low dimensional embedding algorithms, such as Uniform Manifold Approximation and Projection (UMAP)^30^ and t-Distributed Stochastic Neighbor Embedding (t-SNE),^31^ which can capture local neighborhood structure and overcome the linearity assumption of PCA, construct axes which lack interpretability. To address these challenges, we sought to “go beyond the manifold” by combining the predictive power of partial differential equations with dimensional embedding algorithms, aiming to construct an explainable map of human gastruloid morphospace.

Here, we integrate computational modeling with high-throughput experimental techniques to construct a detailed and predictive map of human gastruloid morphospace. By coupling partial differential equation models of BMP, Wnt, and Nodal signaling dynamics with dimensionality reduction algorithms like t-SNE, we systematically explore how various perturbations influence cell fate patterning within the 2D gastruloid model. We identify two key parameters, cell density and SOX2 stability, that define the major axes of pattern variability, and reveal how these factors interact to influence developmental outcomes. We demonstrate that these parameters explain both canonical and non-canonical patterning of the 2D gastruloid, including specific failure modes linked to teratogenic drugs. This framework not only advances our understanding of the molecular mechanisms underlying human gastrulation but also provides a quantitative, human-specific platform for predicting teratogenic risk, identifying potential therapeutic targets, and evaluating developmental toxicity in a high-throughput, scalable manner.

## Results

### A platform for high-throughput perturbations and analysis of the 2D gastruloid

Many questions about human gastrulation remain unanswered due to ethical constraints on large-scale experimentation with human embryos, as well as significant differences in both embryo anatomy and signaling between humans and mouse, our closest mammalian model. We sought to leverage the 2D gastruloid model in a high-throughput screen to map the morphospace of human gastrulation based on prior demonstrations that the model mimics the cell fate decisions of human gastrulation and produces well-defined spatial regions of differentiation that correspond to cell fate specification of the primitive streak (Figure 1A-B). We reasoned that mapping the differentiation outcomes to a wide variety of perturbation would answer the following outstanding questions: (1) What are the failure modes of human gastrulation? (2) What physiochemical variables are required to accurately model germ layer patterning? (3) What developmental parameters explain the variability in patterning outcomes? And finally, (4) can we build a quantitative model that predicts human teratogens?

**Figure 1:**
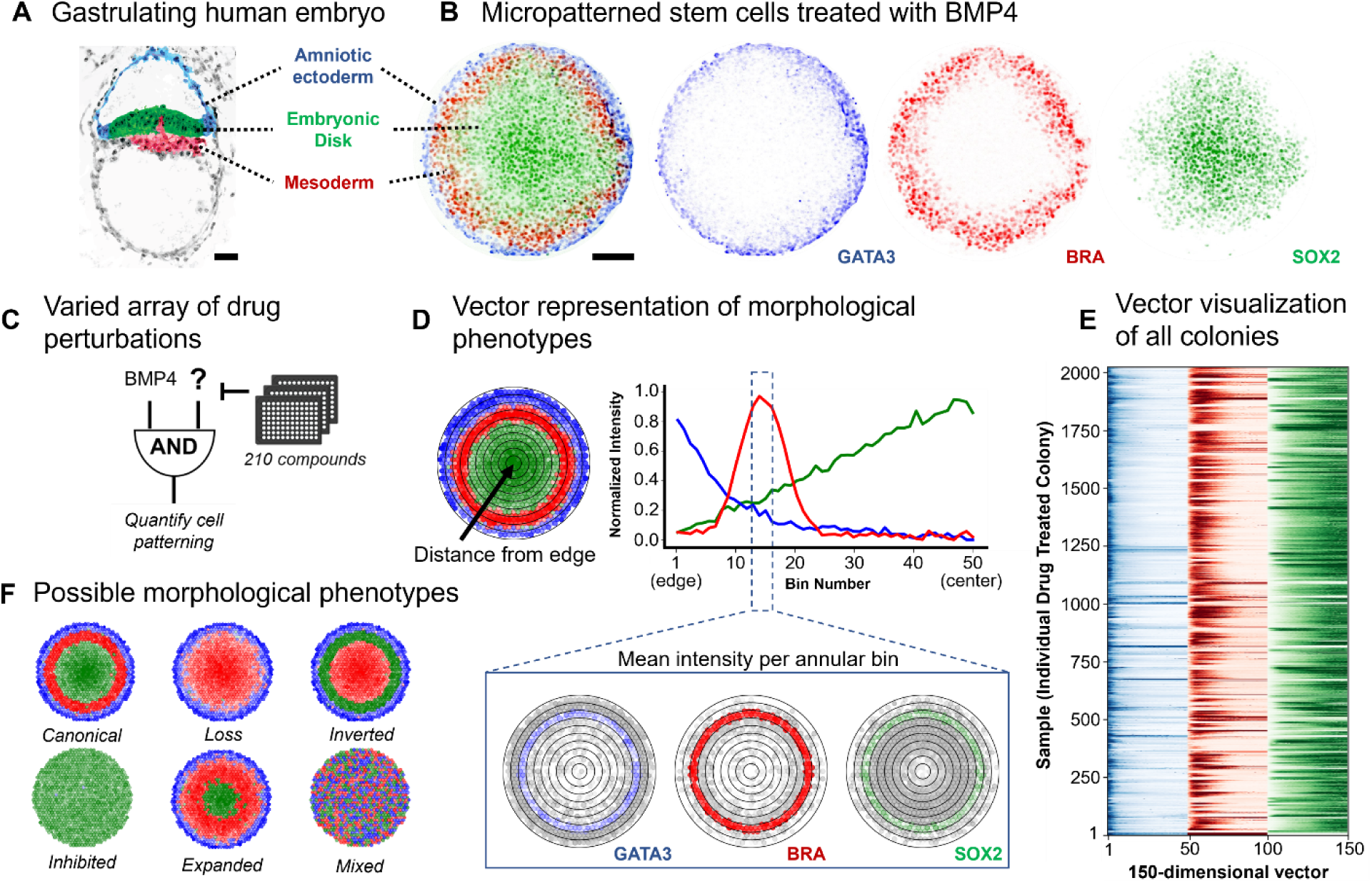
Framework for quantitative characterization and perturbation of morphological phenotypes in 2D gastruloids. (A) Schematic of human embryo showing cell fates observed during gastrulation: amniotic ectoderm (blue), embryonic disk (green), and mesoderm (red). The schematic is based on Carnegie Embryo #7801 from the digitally reproduced embryonic morphology (DREM) project (https://www.ehd.org/virtual-human-embryo/). (B) Representative BMP4-treated 2D gastruloid with immunofluorescence staining for GATA3 (amniotic ectoderm, blue), BRA (mesoderm, red), and SOX2 (embryonic disk, green). (C) Schematic showing the combinatorial drug screening approach used to perturb gastruloid patterning, testing the effects of various drug treatments on cell patterning. (D) Approach for quantifying cell patterning in 2D gastruloids by segmenting radial sections into 50 azimuthal bins from edge to center. The intensity of each bin reflects the mean nuclear intensity of all nuclei located within that radial bin. A sample line plot shows average signal intensity profiles for GATA3, BRA, and SOX2 across the colony radius, demonstrating spatial organization of cell dates. (E) Heatmap representing the 150-dimensional vectorized morphological features of colonies, with bins 1-50, 51-100, and 101-150 representing GATA3, BRA, and SOX2 mean signal intensity from colony edge to center, respectively. (F) Schematic illustrating hypothetical morphological phenotypes, highlighting potential variations in patterning outcomes due to combinatorial perturbations. Scale bars: (A) 50 μm, (B) 100 μm.

We chose to perturb BMP4-initiated gastruloids with compounds from a library of 210 drugs annotated for their activity against stem cell signaling pathways (Figure 1C). While it is possible that a given drug may have multiple protein targets, our approach is target-agnostic and only requires a large diversity of perturbations to capture the variation in outcomes. To quantify phenotypes, we chose to use immunofluorescence staining of germ layer markers: GATA3 for amniotic ectoderm, Brachyury (BRA) for mesoderm, and SOX2 for the undifferentiated embryonic disk. We imaged ∼10 colonies per drug condition and used a custom image segmentation algorithm to identify the levels of each cell fate marker in every nucleus of each colony (Figure S1A), collecting both cell fate and location for ∼2 million cells across 2,025 colonies.

We leveraged the fact that colonies are generally radially-symmetric to build a data structure for downstream pattern analysis. Inspired by approaches that vectorize animal behavior,^32^ we reasoned that we could meaningfully compress colony morphology through averaging cell fates over a set of 50 azimuthal bins, each ∼5 µm in width, ranked by their position, from the edge to the center of each colony (Figure 1D). This yielded a 150-dimensional vector for each colony containing the average azimuthal signal for GATA3, BRA, and SOX2 as a function of colony position (Figure 1E). We hypothesized that this data structure would allow use to easily categorize a wide variety of potential phenotypes, ranging from expansion and retraction, to inversion and even completely mixed orientation of a germ layer fates (Figure 1F). As controls, we included several BMP4-only and untreated (no-BMP4) colonies from multiple independent wells, as well as a Wnt-activating (CHIR-98014) positive control which, as expected, differentiated the entire colony into mesoderm (Figure S1B). These controls demonstrated that patterning phenotypes were relatively reproducible across multiple wells, and days of running the assay (Figure S1C), confirming that the 2D gastruloid model analyzed in this way meets the criteria for constructing a morphospace map.

### Human Gastruloid Morphospace

Upon visual inspection of the drug-treated gastruloids, we noticed a variety of deviations from the canonical patterning observed in the controls. We sought to develop an unsupervised algorithm that would (1) project each colony down to 2 dimensions and (2) identify groups of perturbations that result in similar phenotypes so that they could be evaluated as potential failure modes. Inspired by previous methods,^33^ we coupled t-SNE to an unsupervised clustering procedure which applies watershed segmentation to the two-dimensional embedding after convolution with a gaussian kernel (Figure S2A, Methods). This method does not impose a fixed number of clusters, as other clustering algorithms do (k-means, Gaussian mixture models, spectral clustering, etc.), and thus produces a set of groups that reflect the local variations in phenotype without supervision (Figure 2A). We identified 12 clusters (C1-C12) which we call morphological phenotypes.

**Figure 2:**
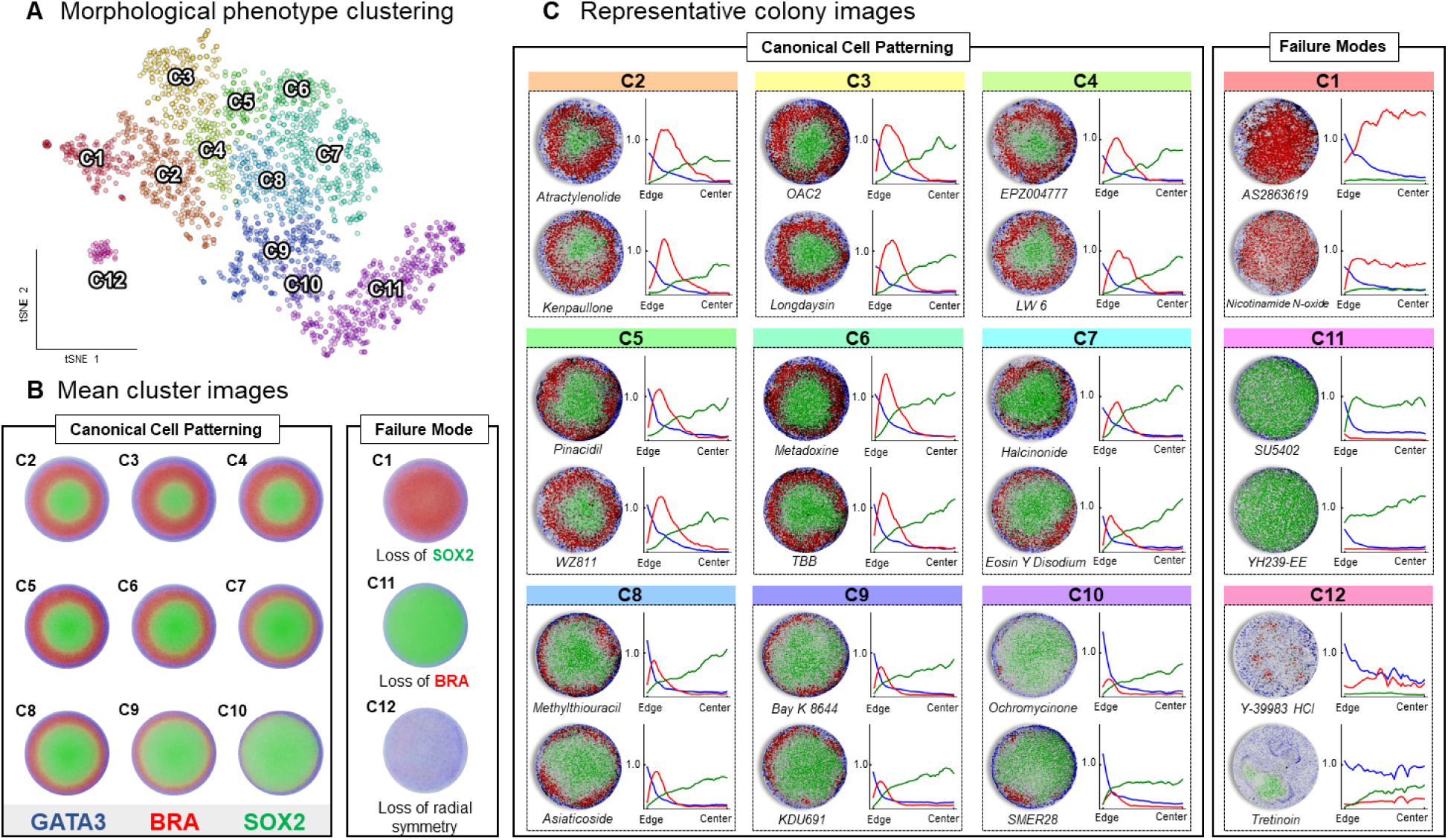
Mapping the morphospace of 2D gastruloids to identify human gastrulation failure modes. (A) t-SNE plot of individual 2D gastruloids, each represented as a point and clustered by morphological phenotype based on high-dimensional image analysis. An unsupervised clustering algorithm was applied to identify distinct clusters, resulting in the identification of 12 unique morphological phenotypes. (B) Mean cluster images for each morphological phenotype, generated by averaging all colony images within a cluster. This approach highlights essential features of canonical cell patterning and failure modes, facilitating the identification of key morphological differences, such as loss of SOX2, loss of BRA, or loss of radial symmetry. (C) Representative colony images and associated drug treatments for each of the 12 clusters, accompanied by radial line plots showing the spatial distribution of cell fate markers (GATA3, BRA, SOX2) from edge to center of the colonies. The distinct and recurring patterns observed within each cluster underscore the effectiveness of this framework in defining and categorizing morphological phenotypes, revealing how various perturbations impact patterning in the 2D gastruloid.

To visualize the phenotype of each cluster, we generated composite images by taking the mean pixel intensity for each cell fate marker from all colonies in each cluster (Figure 2B). This allowed us to build a coarse-grained understanding of colony patterning across the entirety of morphospace. Clusters could be divided into two major categories. The first group, describing the majority of colonies (C2-C10), displayed changes in the relative size of each region marked by cell-fate markers but did not alter the basic topology of the patterns. We termed this central region of morphospace the canonically patterned region. The second group, colonies outside of this region (clusters C1, C11, and C12), displayed either the loss of a cell-fate marker and/or loss of radial symmetry (Figure 2B-C). Since either of the aberrations captured in these clusters would result in a failure of gastrulation, we termed each of these clusters to be gastrulation failure modes. For more examples see Figure S2B-C and for the entire dataset visit https://max-wilson.mcdb.ucsb.edu/research/gastruloid-morphospace.

### Cell density explains mesoderm positioning and rescues malformation-inducing drugs

We next turned our attention to determine the causes of variance in the canonical patterning region. We observed that one of the most obvious sources in this region is the width of the mesoderm band (Figure S3A-B). From previous observations on the importance of 2D gastruloid density in determining patterns,^28,34–36^ we hypothesized that mesoderm band width could be regulated by cell density. This implies that a significant number of perturbations influence pattern formation by simply altering the rates of cell division or death during the 48-hour differentiation. Thus, we first examined experimental fluctuations in cell density.

The number of cells that attach to each micropattern is Poisson distributed, thereby providing a natural variation in the cell density parameter. This allowed us to test the cell density hypothesis in our BMP4-treated controls. Indeed, the distribution of these control colonies in morphospace is explained by their density (Figure 3A). We also observed a correlation (R^2^ = 0.69) between cell density and the location of the peak of the mesoderm band (Figure 3A). Finally, ranking the edge-to-center mesoderm staining for each BMP4 control colony by density also revealed a clear trend of mesoderm expansion as cell density decreased (Figure 3B). Other drug treatments, targeting a variety of signaling pathways, also displayed this relationship at the single-colony level (Figure S3C-I).

**Figure 3:**
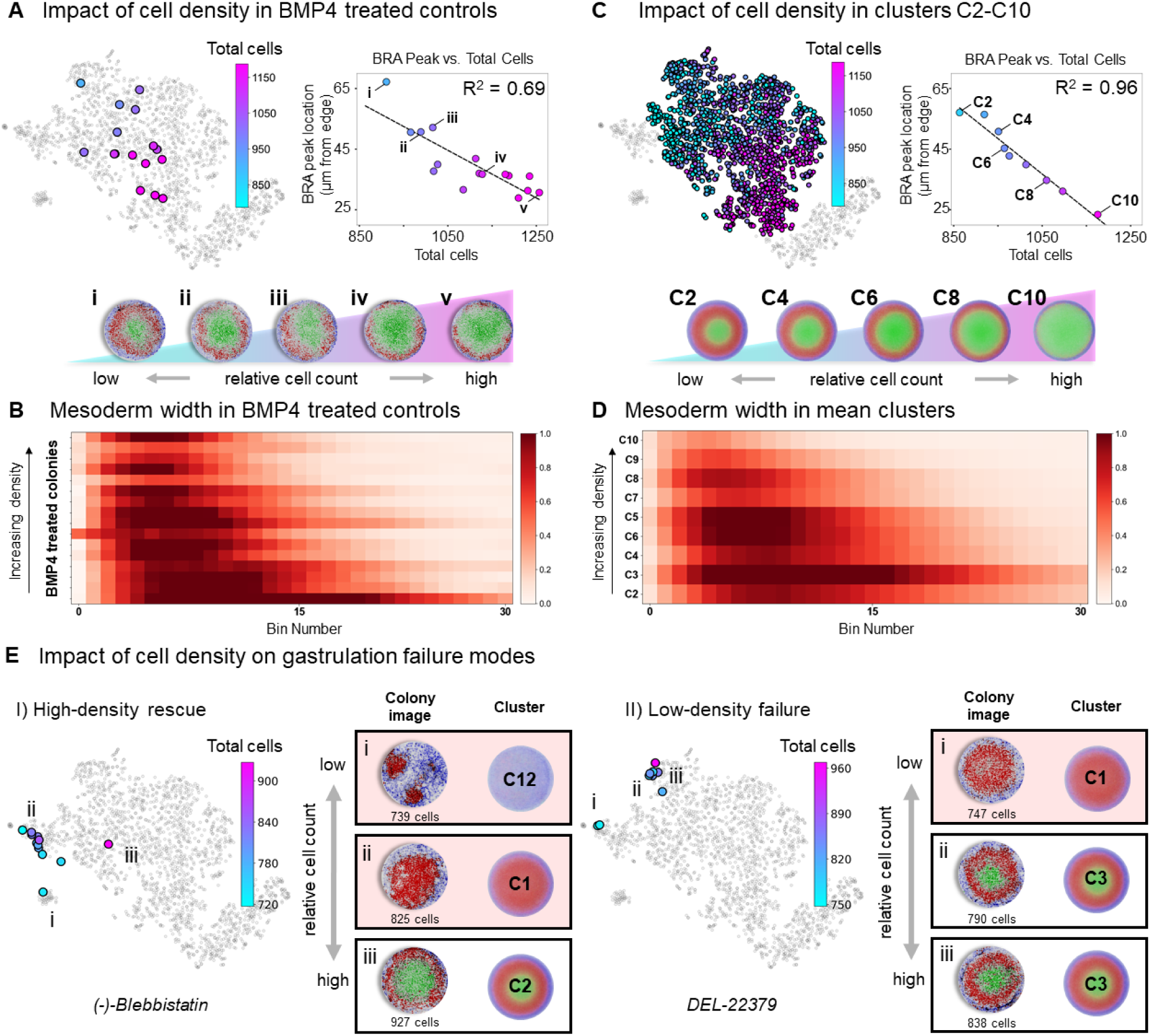
Impact of cell density on mesoderm patterning and rescue of drug-induced failure modes in 2D gastruloids. (A) Analysis of cell density effects on patterning in BMP4-treated control colonies. Colonies are plotted in morphospace and colored by cell density, with the inset showing a strong correlation (R² = 0.69) between cell density and the inward shift of the BRA (mesoderm) peak. Representative images of highlighted colonies illustrate a clear trend of decreasing mesoderm band thickness with increasing cell density. (B) Heatmap showing edge-to-center mesoderm staining intensity for BMP4-treated controls, ranked by cell density. Results reveal mesoderm expansion and inward shift at lower densities. (C) Clusters C2-C10 are shown in morphospace, colored by cell density, with the inset revealing a significant correlation (R² = 0.96) between mesoderm peak location and cell density in mean cluster representations. Images of mean clusters further confirm that mesoderm location and width are influenced by cell density across the canonical patterning region. (D) Heatmap of mesoderm width in mean cluster representations of C2-C10, demonstrating that lower cell densities are associated with broader mesoderm bands across the canonical patterning region. (E) The impact of cell density on rescuing or exacerbating gastrulation failure modes. (I) High-density conditions can rescue colonies from failure mode clusters (e.g., (-)-Blebbistatin-treated colonies from C1 and C12 are driven back into canonical patterns). (II) Conversely, low-density conditions can push drug-treated colonies (e.g., DEL-22379) into failure modes. These results suggest that cell density plays a significant role in modulating patterning outcomes and can potentially correct aberrant gastrulation phenotypes.

To determine if this relationship was also true across all colonies in the canonical patterning region, we performed a similar analysis on the composite colonies for each cluster. Again, we observed a clear relationship between cell density and a colony’s location in morphospace and an even more significant correlation (R^2^ = 0.96) between the mean cluster density and the location of the mesoderm band peak when plotted by cluster (Figure 3C). Finally, we again confirmed this relationship by ranking the mean edge-to-center mesoderm vectors for each cluster by their mean cell density (Figure 3D). Thus, both within single-drug treatments and across all canonically patterned phenotype clusters a major source of variance is explained by cell density. Ultimately, density is regulated by the balance between cell division and death, processes which incorporate information from many parallel pathways and thus encompass a large number of drug targets.

Given the impact of cell density on patterning, we hypothesized that it might be strong enough to override the effects of drug treatment. If correct, we would expect colonies to be rescued back into the canonical patterning region by increased density or driven out into failure modes by decreased density, demonstrating the capacity of cell density to modulate drug-induced patterning outcomes. To this end, we observed (-)-Blebbistatin treated colonies, which were typically in failure mode clusters C1 and C12, that could be rescued through increasing the cell density (Figure 3E I). We also observed colonies driven to the edge of the canonical patterning region by a drug treatment (e.g. DEL-22379 and additional examples in Figure S3J) drop-out of the canonical patterning region when their density becomes too low (Figure 3E II). Overall, these observations suggest that density is causal of a large amount of variation in 2D gastruloid patterning and could be used as a target for correcting aberrant gastrulation.

### A mathematical model of density-induced variations in morphogen signaling dynamics explains mesoderm band variation

Various studies have developed models of the effects of individual morphogens on cell fates.^37–39^ While combinatorial morphogen approaches have recently been applied to model cell fate decisions in non-patterned stem cells,^40^ as well as the 2D gastruloid,^41^ we lack a comprehensive model that incorporates combinatorial signaling dynamics and how those dynamics are decoded into cell fates. Such a model would enable an unbiased analysis of patterning outcomes in the gastruloid based on their fate patterning. Building off of previous partial differential equation models of morphogen signaling,^41^ we utilized a reaction-diffusion description of BMP, Wnt, and Nodal dynamics. To model the fate choices, we further incorporated equations that describe how the morphogens are decoded into each of the three cell fate markers, GATA3, BRA, and SOX2 (Figure 4A-B, Figure S4A, and Methods). We note that existing models of cell fate in the 2D gastruloid incorporate entire signaling histories, but only focus on a single morphogen,^37,40^ or incorporate multiple morphogens, but neglect dynamics and make heuristic fate predictions based only on instantaneous morphogen levels.^41^

**Figure 4:**
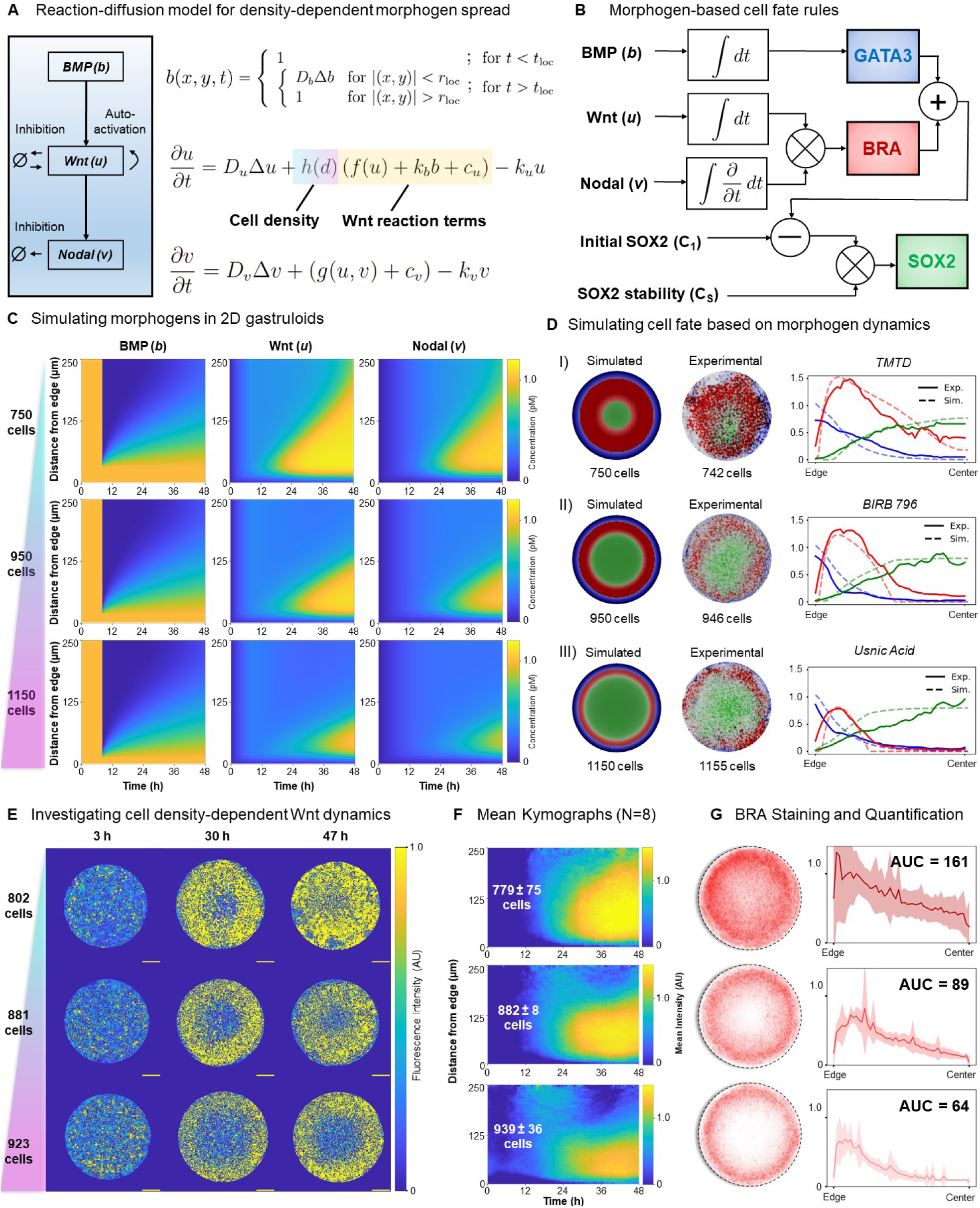
Modeling the impact of cell density on morphogen dynamics and cell fate determination in 2D gastruloids. (A) Reaction-diffusion model describing the density-dependent spread of morphogens BMP, Wnt, and Nodal in 2D gastruloids. The model incorporates cell density effects by modifying Wnt reaction terms. (B) Logic-based schematic diagram of cell fate determination rules based on integrated morphogen dynamics. Notably, cells respond to the integral of BMP and Wnt (concentration-dependent), but to the integral of the partial derivative of Nodal, or the rate of Nodal concentration change (rate-dependent). Initial SOX2 levels (C_1_) and SOX2 stability (C_s_) modulate the response of cells to these signals, influencing differentiation outcomes. (C) Simulated concentration profiles of BMP, Wnt, and Nodal over time in 2D gastruloids at varying cell densities (750, 950, and 1150 cells). The simulations reveal that lower cell densities lead to faster and deeper penetration of Wnt and Nodal signals into the colony, influencing mesoderm patterning. (D) Simulated versus experimental cell fate outcomes for different densities, demonstrating the model’s ability to replicate patterns observed in our screen. Three examples of simulations (left) compared to experimental images (right) with accompanying line plots show matching spatial distributions of cell fate markers (GATA3, BRA, SOX2). (E) Experimental investigation of density-dependent Wnt dynamics using CRISPR-tagged β-catenin (β-cat) hESC lines. Live-cell time-lapse imaging over 48 hours shows the effect of cell density on Wnt signaling. Scale bars = 100 μm. (F) Mean kymographs of Wnt signaling activity across 8 colonies per cell density range, demonstrating that lower cell densities result in a faster and broader spread of the Wnt signal. (G) Quantification of BRA staining intensity profiles across different densities, taken from the same colonies used to generate the kymographs. The shapes of the curves show that BRA signal penetrates further inward as cell density decreases, and the area under the curve (AUC) analysis confirms increased total BRA signal at lower densities, validating our model’s prediction of density-induced variations in mesoderm dynamics.

Based on our observations of the importance of cell density, we sought to include its effects in our model as well. Because hESCs form a tight monolayer, but also cannot grow beyond the bounds of the micropatterned ECM onto which they are plated, multiple morpho-mechanical features of cells in the gastruloid change as a function of density (e.g. aspect ratio, cortical tension, receptor availability, volume, etc.). We hypothesized that Wnt signal secretion, with its reported sensitivity to changes in local mechanics,^42^ is regulated by cell density. Mathematically, we captured this by introducing a function, h(d), that linearly decreases all sources of Wnt ligand creation as density, *d*, increases. For details of model implementation and further justification of parameter choices see Methods.

To test if we accurately captured the effects of density, we simulated morphogen dynamics and cell fate determination of three colonies that range in densities, from 750 cells (∼10^th^ percentile) to 1150 cells (∼90^th^ percentile). Kymographs of simulated signaling revealed a faster Wnt wave that further permeates the colony interior as density decreases (Figure 4C). As a result, Nodal, which is created as a function of Wnt, also follows this trend. Ultimately, this leads to a deeper penetration of BRA into the center of the colony with a corresponding decrease in SOX2 because BRA is a function of both Wnt and Nodal (Figure 4D, Video S1). Experimental colonies with similar densities matched simulation results across a wide range of density, even at the extremes of density and complete inhibition of a morphogen signal (Figure S4B-C). These findings suggest that our comprehensive model accurately captures the interactions between morphogen signaling, cell density, and cell fate markers.

A central implication of our model is that Wnt signaling is inversely proportional to cell density. To test this prediction, we sought to experimentally measure endogenous Wnt signaling in 2D gastruloids as a function of cell density. We leveraged a CRISPR-tagged β-Catenin (β-cat)^43^ hESC line (Methods) to observe Wnt signaling in real time. We varied initial seeding density and took live-cell, time-lapse images of 8 colonies per condition over the 48 hr differentiation (Figure 4E). To extract the β-cat signaling activity we wrote a custom image processing algorithm that extracts the active, non-membrane β-cat from the inactive, membrane-bound pool, as has been done previously^41^ (Figure 4SD). Both timelapse videos from individual colonies (Figure S4E-G, Video S2) and mean kymographs, subset by colony density, clearly demonstrate the impact of cell density on the Wnt wave (Figure 4F). Finally, we fixed and stained the same colonies for which β-cat was imaged and confirmed that their BRA staining also decreased with increasing density (Figure 4G). Overall, these experiments validate our choice of model architecture, confirm the effects of cell density on Wnt signaling, and suggest that our cell fate rules accurately capture the decoding process that occurs in the gastruloid.

### Cell density and SOX2 stability explain variance in canonical patterning

The non-parametric approach t-SNE and related dimensionality reduction techniques employ to generate embeddings suffer from two important limitations: (1) the lack of interpretable axes in the low-dimensional space and (2) the dependence of the embedding on the entire dataset, which precludes the addition of new data. We reasoned that overcoming these limitations would allow us to place simulated colonies into the experimentally-defined morphospace, thereby revealing the relationship between developmental parameters and location in morphospace. This would have the added benefit of inferring drug mechanisms by mapping changes in model parameters onto locations in morphospace where specific drugs reside.

While there is no inverse solution to t-SNE, one can construct a function, 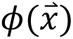, that approximates the low-dimensional embedding. To construct 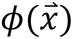 we trained a 4-layer neural network with ReLu activation functions to minimize the distance between the learned location in the low-dimensional embedding and the true location from our original t-SNE (Figure 5A, see Methods for details). The learned embedding was able to accurately reconstruct the original morphospace map (Figure S5A). To understand the universality of the density effect on patterning, we used 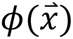 to project simulated colonies with varied density, *d*, and all other parameters kept constant (Figure 5B). Intriguingly, these simulated colonies formed a smooth and continuous curve, suggesting that this single parameter describes one of the major axes of variance over the canonical patterning region (Figure 5C).

**Figure 5:**
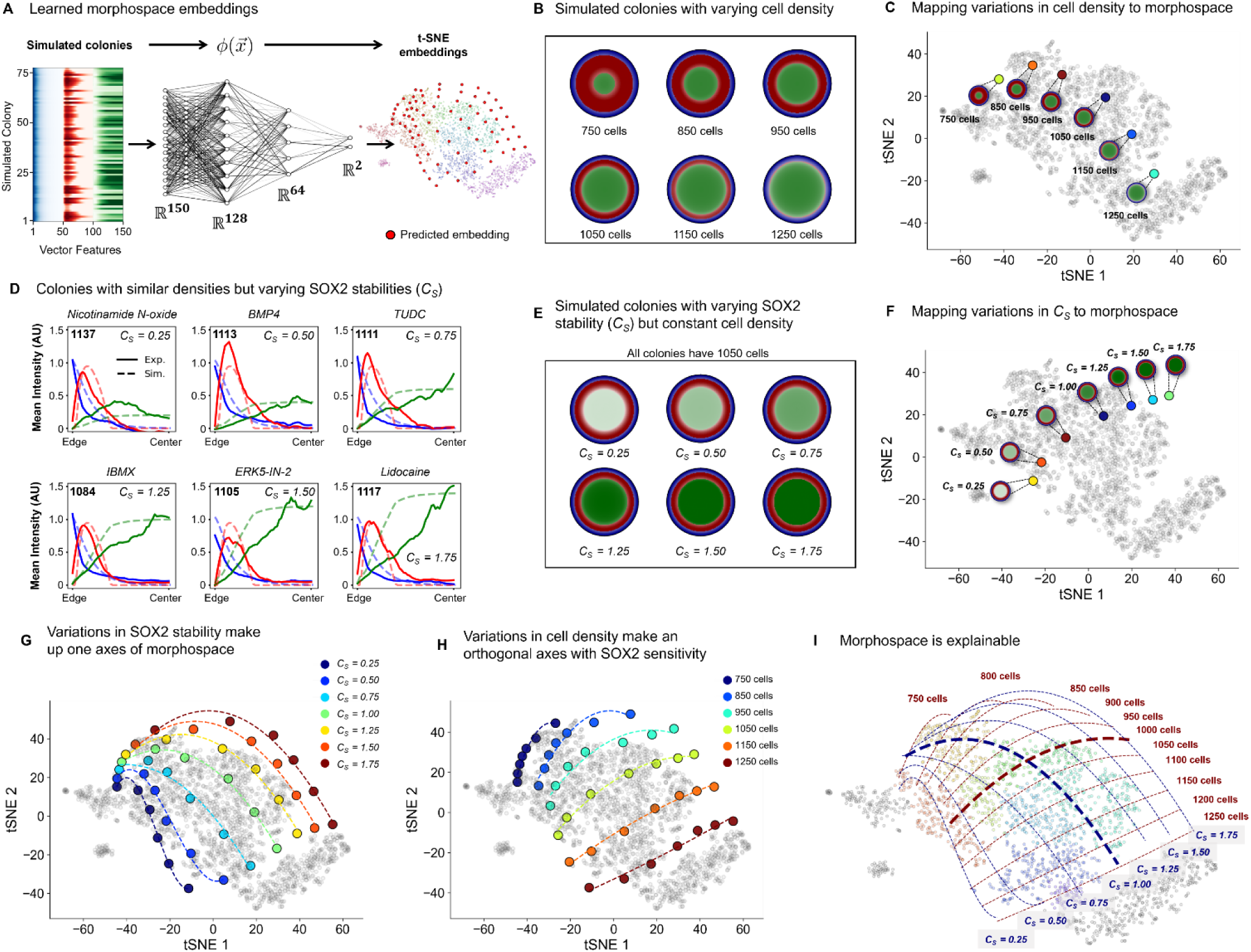
The Morphospace of 2D Gastruloids is Explained by Two Parameters: Cell Density and SOX2 Stability. (A) Schematic showing the projection of simulated colonies into the experimentally defined morphospace using a neural network-based function, ϕ(x), trained to map high-dimensional data into the low-dimensional t-SNE space. (B) Simulated colonies with varying cell densities, showing how simulated spatial patterns align with the experimental trend of decreasing mesoderm thickness as cell density increases. (C) Representative simulated colonies projected onto the t-SNE embedding of experimental data, matching the expected locations in morphospace. (D) Simulated versus experimental intensity profiles of cell fate markers across a range of SOX2 stability values (C_s_), with simulated density kept constant at 1100 cells. The profiles demonstrate how changes in C_s_ affect patterning, with experimental cell density values shown at the top left of each plot, within ±5% of the simulated density. (E) Simulated colonies with varying SOX2 stability, but constant cell density. SOX2 stability influences the final expression of SOX2 relative to GATA3 and BRA. (F) Projection of simulated colonies with varying SOX2 stability values into morphospace, forming a curve that appears orthogonal to the cell density axis, suggesting SOX2 stability is a distinct driver of pattern variability. (G-H) Combined plot showing simulated colonies projected along variations in both cell density and SOX2 stability, forming a grid that captures the full range of canonical patterning outcomes observed experimentally. (I) Comprehensive map of the canonical patterning region, with fitted polynomials forming a grid that defines putative axes of morphospace, enabling precise description of a colony’s location within morphospace. This approach demonstrates that manipulating just these two parameters—cell density and SOX2 stability—recapitulates the majority of observed patterning outcomes, making the morphospace interpretable and highlighting the key developmental parameters driving pattern variability.

In examining the experimental variation in colonies with similar densities we noticed that an independent axis of variation might be described by the shape and level of SOX2 (Figure 5D). Thus, we hypothesized that the SOX2 stability, represented by the *C_s_* parameter in our simulation, forms a second axis. We therefore simulated colonies varying only *C_s_* and, after projecting them into morphospace, found that they indeed form a second seemingly orthogonal axis (Figure 5E-F). To understand if these two parameters alone could explain the entire canonical patterning region, we systematically varied *C*_s_ and *d* (Figure S5B). Projecting all simulating colonies into morphospace revealed a grid (Figure 5G-H). Fitting polynomials to these simulated colonies demonstrated that the entire canonical patterning region is explained by varying just two parameters, *C_s_* and *d* (Figure 5I). Thus, our framework makes the 150-dimensional manifold interpretable and describes 72 percent of all patterning outcomes. Of note, while t-SNE is sensitive to the hyperparameters chosen for the embedding, the impact of cell density on patterning, as well as the *C_s_* and *d* axes are maintained even when choosing an alternative seed for the t-SNE embedding, indicating the robustness and consistency of our approach in capturing key drivers of pattern variability, independent of embedding conditions (Figure S5C-H).

### Non-canonical gastruloid patterns represent unique failure modes and predict teratogens

Having explained the canonical patterning region, we turned our attention to colonies that fall outside this part of morphospace: clusters C1, C11, and C12 (Figure 6A). As mentioned above, cluster C1 is characterized by BRA overexpression. We noticed that drugs inducing this phenotype tend to cause tonic activation of the Wnt pathway (e.g., CHIR-99021, CHIR-99014, etc.), inhibit Rho Kinase, and generally appear to be useful for increasing regeneration (PDGH-1 inhibitor) (Figure 6B). As Rho-kinase is known to alter the cell’s mechanics and perception of its neighborhood,^44^ this further confirms the relationship between mesoderm patterning and cell density. Cluster C12 is characterized by low cell number, high GATA3, and a general loss of radial symmetry (Figure 6C). Drugs in this cluster appear to be toxic to hESCs, inhibit mTOR (Rapamycin), or induce severe malformations (Tretinoin). Protocols designed to differentiate amnion-like cells may benefit from drugs in this cluster.

**Figure 6:**
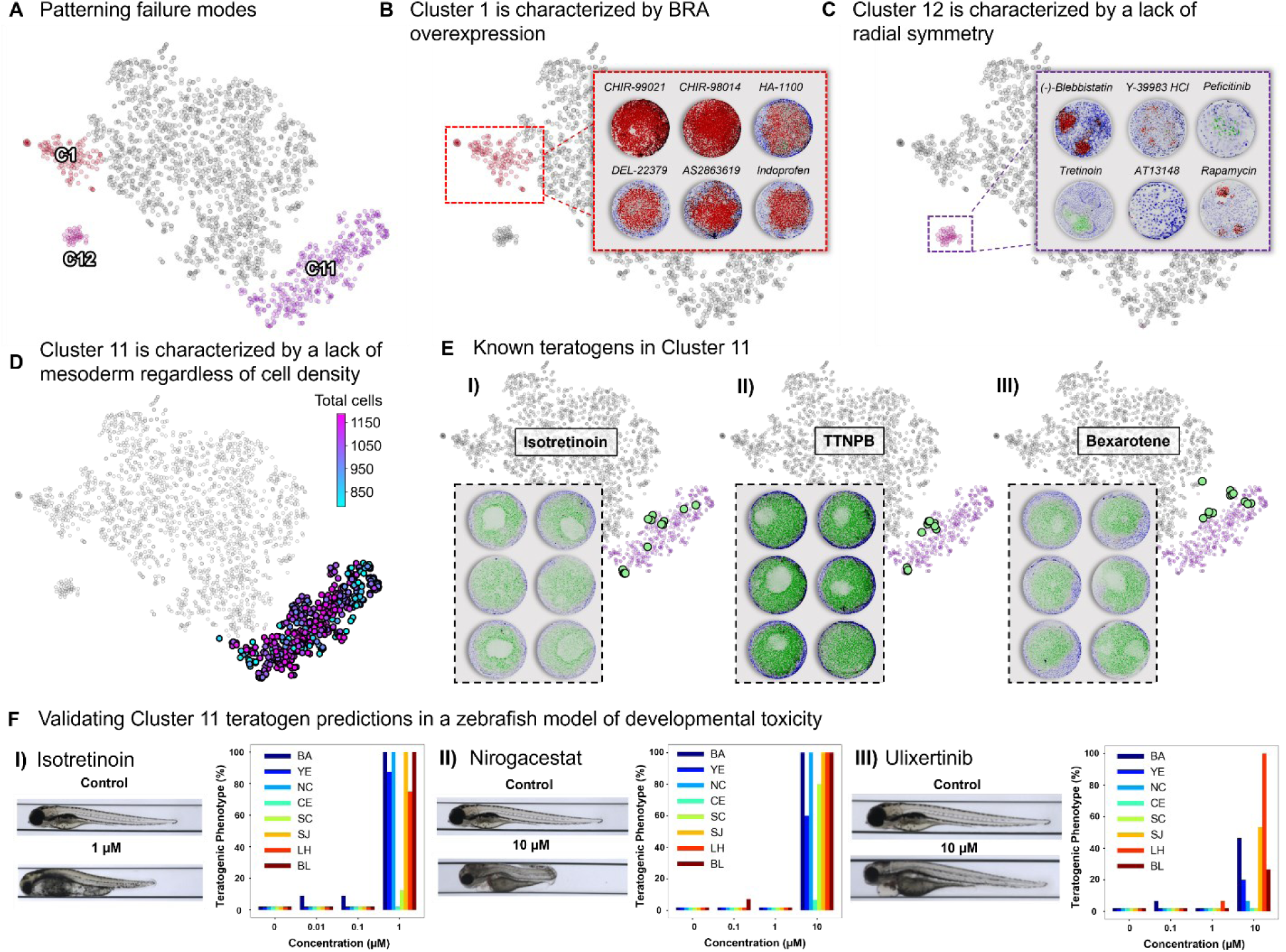
Identification and Characterization of Non-Canonical Patterning Clusters in 2D Gastruloids as Failure Modes of Human Gastrulation. (A) Clusters C1, C11, and C12 are identified as non-canonical patterning regions within the 2D gastruloid morphospace, representing distinct failure modes of gastrulation characterized by specific morphological anomalies such as loss of SOX2, overexpression of BRA, and loss of radial symmetry, respectively. (B) Cluster C1 is defined by colonies with BRA overexpression, often induced by Wnt pathway agonists (e.g., CHIR-99021), highlighting a failure mode linked to excessive Wnt signaling. (C) Cluster C12 is characterized by colonies displaying low cell numbers, loss of radial symmetry, and high levels of GATA3, commonly caused by toxic compounds or drugs targeting Rho-kinase and mTOR (e.g., (-)-Blebbistatin, Y-39983 HCl, Rapamycin). (D) Cluster C11 exhibits a failure mode marked by the loss of BRA and disrupted cell patterning, regardless of cell density. (E) Known teratogens identified in Cluster C11, including Isotretinoin, TTNPB, and Bexarotene, show severe disruptions in gastruloid patterning, indicating high teratogenic risk and corresponding failure modes of gastrulation. (F) Zebrafish assays were used to validate the teratogenic potential of drugs identified in Cluster C11. Isotretinoin, a known teratogen, was selected as a positive control, alongside two other drugs from Cluster C11—Nirogacestat and Ulixertinib—that are previously unclassified as teratogens. Treated embryos exhibited qualitative and quantitative defects consistent with failure modes in embryonic development, including abnormal body curvature (BA), yolk edema (YE), necrosis (NC), craniofacial edema (CE), scoliosis (SC), snout and jaw defects (SJ), reduced lateral heart area (LH), and decreased lateral body length (BL), as indicated by the color-coded bars.

Of all the non-canonical clusters, C11 has by far the largest number of colonies (20% of all colonies). Interestingly, colonies falling in this cluster appear to have lost the relationship between *d* and mesoderm, indicating a change to the underlying molecular topology (Figure 6D). Confirming this, we found known inhibitors of BMP, Wnt, and Nodal. In addition, we found a number of known severe teratogens^45–47^ that do not directly inhibit signaling pathways, but are instead thought to activate developmental pathways such as retinoic acid signaling (Figure 6E). Retinoic acid derivatives in C11 also cause 3D protrusion from the 2D gastruloids. Thus C11, appears to describe a failure mode that is non-cytotoxic and uncouples developmental signaling pathways from establishing mesoderm, thereby describing perturbations that fundamentally alter the molecular relationships that determine cell fate.

We noted many drugs in C11 that have not been formally classified as teratogenic and wondered if morphological clustering could provide a method to assess teratogenic risk. To this end we selected one well known teratogen, Isotretinoin, and two clinical-stage molecules of unknown teratogenic risk, Nirogacestat and Ulixertinib, to test in a zebrafish-based assay of developmental teratogenicity. Analysis of zebrafish developmental outcomes has been shown to have 74.19% accuracy (87.50% sensitivity) in comparison to mice at 67.74% (75.00% sensitivity),^48^ and are also faster and easier to test at scale. Examination of gross anatomy as well as specific qualitative and quantitative teratogenic phenotypes revealed that all molecules tested are highly teratogenic (Figure 6F). Overall, these findings demonstrate that morphological analysis of the 2D gastruloid could be a highly accurate and quantitative assay to evaluate potential teratogenic risk of a compound, especially during gastrulation–the most susceptible window of embryonic development.^49^

## Discussion

Our understanding of morphogen signaling has evolved from a static picture of standing gradients, such as the anterior-posterior-determining Bicoid gradient observed in the *Drosophila*, to a dynamic picture involving waves.^41,50^ Concurrently, models that predict cell fate from morphogen signaling have progressed from static threshold measuring mechanisms to dynamic processes such as integral decoding.^37,51^ Meanwhile, our ability to map the state-space of outcomes resulting from these complex, dynamical processes has also progressed by coupling high-throughput, high-dimensional measurement techniques to manifold embedding and dimensionality-reduction algorithms that enable the visualization of high-dimensional data. Here, we applied these techniques to comprehensively understand how morphogen dynamics determine developmental outcomes in the 2D gastruloid. Our low-dimensional mapping, or morphospace, enables the visualization and comparison of patterning outcomes as well as the identification of failure modes of human gastrulation.

### Going beyond the manifold

While non-linear dimensionality techniques are an excellent approach for identifying trends and clusters from high dimensional data, which in the context of the 2D gastruloid can be considered phenotypes, there are no existing methods for linking the partial differential equations that describe the relationship between state variables to their outcomes. As a result, non-linear dimensionality techniques lack inherent explainability– it is impossible to know which system parameters determine the location of a point in the low-dimensional embedding. We overcame this limitation by learning the embedding that maps high-dimensional observations to the low-dimensional space. This approach allowed us to explain the low-dimensional embedding and establish the parameters that describe the axes of this space. This framework may be helpful in applying models to describe other biological state spaces, such as those derived from single-cell transcriptomics and high-throughput proteomics.

Leveraging this new approach, we identified cell density and SOX2 stability as two key parameters describing the canonical patterning region, which begs the question: why these two parameters? Cell density is a parameter that is functionally linked to multiple mechanical and chemical signaling pathways, and thereby may serve as a proxy for a variety of environmental cues, including number of neighbors, proliferation capacity, and tissue stiffness. Importantly, a notable difference between the 2D gastruloid and the embryo is that, in the gastruloid, cells are confined to grow on micropatterns. This constraint causes cells to spread at low densities and crowd at high densities, leading to alterations in mechanical properties known to influence patterning in the 2D gastruloid, including membrane tension^52^ and receptor localization.^34^ Thus, the coupling of germ-layer fates to the engineered constraint of cell patterning may increase the sensitivity of this system to detecting perturbations.

Our observations revealed increased Wnt signaling and enhanced mesoderm specification at low cell densities, suggesting that Wnt signaling may have evolved to detect cell density in the embryo through either a chemi- or mechano-sensitive mechanism. Supporting this idea, nuclear translocation of β-catenin can be triggered by mechanical forces and induce mesoderm differentiation in species ranging from zebrafish and *Drosophila*^53^ to the sea anemone.^54^ Together with our findings, these observations suggest an ancient, evolutionary conserved role for density-sensing via Wnt signaling that dates back at least 600 million years.

The orthogonal parameter in morphospace to cell density is SOX2 stability. Similar to cell density, we speculate that it could also serve as a proxy for multiple signaling pathways, that can be varied independently of cell density. Typically, as colonies begin to express BRA, there is a concurrent decrease in SOX2 expression. However, altering the SOX2 stability parameter disrupts this balance. At high SOX2 stability values, cells maintain high levels of both BRA and SOX2. Interestingly, the methylation and phosphorylation state of SOX2, which regulates its degradation and activity,^55^ provides a potential mechanism linking SOX2 stability to broader cellular contexts. Indeed, drugs that decrease SOX2 stability (Table S1) include histone methyltransferase inhibitors, which can alter chromatin structure and affect gene expression, and cyclic AMP inhibitors, which serve as proxies for overall cell metabolism. Similarly, drugs that increase SOX2 stability (Table S2) include histone deacetylase inhibitors, which support pluripotency by enhancing chromatin accessibility. Overall, SOX2 stability appears to capture key aspects of the epigenome and cellular fitness, suggesting a complex landscape over which hESCs can modulate their pluripotency in response to cell health and the environment.

### Cell density sensing in human embryos

Our observations revealed a strong reliance on cell density for patterning outcomes, with even minor fluctuations in initial seeding density being sufficient to shift from canonical patterning to failure modes. This raises the question of how precise patterning is achieved in the human embryo given the potential for cell density fluctuations. Interestingly, whole-embryo imaging of mouse gastrulation has revealed that density variance is globally minimized at the primitive streak,^56^ suggesting the existence of a conserved module that can sense and regulate the local density of cells.

Studies on mouse embryos have also shown that the initiation and progression of gastrulation are closely linked to reaching a threshold number of epiblast cells.^57,58^ Embryos with reduced cell numbers due to impaired proliferation experience delayed gastrulation, indicating the necessity of achieving a minimum density for proper developmental timing. Conversely, double-sized embryos, formed by aggregating two 8-cell stage eggs, attain the critical cell number earlier but still initiate gastrulation at the same time as normal-sized embryos,^59^ hinting at the existence of a density threshold rather than just cell count for triggering gastrulation. These findings suggest that a density-sensing mechanism operates to integrate local cell density with developmental cues, maintaining the spatial and temporal precision essential for proper patterning. While it remains to be demonstrated that aberrant density sensing is responsible for failures of human gastrulation, our study of the 2D gastruloid suggests that this module can respond to a wide range of fluctuations and modulate germ layer fate patterning accordingly, at least in part through changes in Wnt signaling.

### A quantitative framework for teratogen screening

Our successful identification of teratogens from cluster C11 serves as a proof-of-concept demonstration that stem cell-based models of human embryonic development can accurately identify novel teratogenic compounds. Small animal models suffer from species-specific differences, are not scalable, and often fail to accurately identify teratogens due to the challenge of administering doses that induce birth defects without being lethal to the embryo or mother.^60^ As a result, more than 80% of FDA approved drugs lack sufficient data to determine fetal risks,^61^ and an estimated 1 in 16 pregnancies are exposed to teratogenic drugs.^62^ Although previous attempts have been made to use micropatterned hESCs for teratogen testing,^63,64^ the incomplete understanding of how system parameters influence patterning outcomes has hindered the ability to accurately distinguish the impact of drugs from other variables. The high-throughput nature of our technique makes it highly suitable for scaling the evaluation of environmental hazards and pharmaceutical drugs in a cost-effective and efficient manner—an essential need given that the USA alone produces an average of 1,500 new substances each year,^65^ contributing to an estimated 140,000 chemicals that have been synthesized globally to date,^66^ many of which are toxic in small doses and are infeasible to test using animal models.

### Limitations of the study

This framework serves as a critical starting point for identifying potential failure modes of human gastrulation and offers a means to uncover how specific perturbations impact development. However, key signaling pathways, notably FGF signaling,^67,68^ and detailed mechanistic insights remain underexplored. While the framework shows potential for teratogen testing, it was not originally designed for this purpose, and comprehensive quantification of sensitivity and specificity has not been performed. Preliminary data using zebrafish embryos indicate promise, yet known teratogens like valproic acid, which primarily affect neural tube development post-gastrulation,^69^ were not flagged as teratogenic in our assay. Additionally, all compounds were tested at a single active concentration (10 µM), which may underestimate failure modes occurring at higher doses. Future work will involve standardizing protocols, testing across a broader range of concentrations, and expanding the set of known teratogens. In addition, measuring just 3 transcription factors limited our ability to characterize certain cell fate decisions of rare cell types, including primordial germ cells, that emerge during gastrulation.^70^ To characterize these rare, but important events, we plan to integrate additional protein markers and small molecule probes, such spatial transcriptomics techniques.^71^ Despite these limitations, our study provides a foundation for future investigations aimed at refining teratogen detection and enhancing our understanding of the molecular underpinnings of human gastrulation.

## Resource Availability

### Lead Contact

Further information and requests for resources and reagents should be directed to and will be fulfilled by the lead contact, Maxwell Wilson.

### Materials Availability

Plasmids generated in this study are available upon request to Maxwell Wilson without restrictions.

### Data and Code Availability

All code generated during this study has been deposited at the Wilson Github repository (https://github.com/mzwlab). An interactive dataset is also available at: https://max-wilson.mcdb.ucsb.edu/research/gastruloid-morphospace. Any additional information required to reanalyze the data reported in this paper is available from the lead contact upon request.

## Acknowledgements

This study was supported by UCSB start-up funds. J.R. is supported by NICHD of the National Institutes of Health via a Ruth L. Kirschstein Postdoctoral Individual National Research Service Award (1F32HD114539-01). The authors acknowledge the use of the Microfluidics Laboratory within the California NanoSystems Institute, supported by the University of California, Santa Barbara and the University of California, Office of the President. Results related to the zebrafish teratogenicity assay were performed in collaboration with ZeClinics. The authors also thank Angela Pitenis and Dennis Clegg for their valuable feedback and discussions on experimental and figure design.

## Author contributions

Conceptualization, J.R., C.Q., D.H., and M.Z.W.; methodology, J.R., C.Q., D.H., N.B., G.D., and M.Z.W.; software and data analysis, J.R., C.Q., and M.Z.W.; modeling and simulations, J.R. and M.Z.W.; investigation, J.R., C.Q., D.H., N.B., G.D., and M.Z.W.; writing – original draft, J.R. and M.Z.W.; editing – final draft, J.R. and M.Z.W.; funding acquisition, J.R. and M.Z.W.

## Supplemental Figures

**Figure S1:**
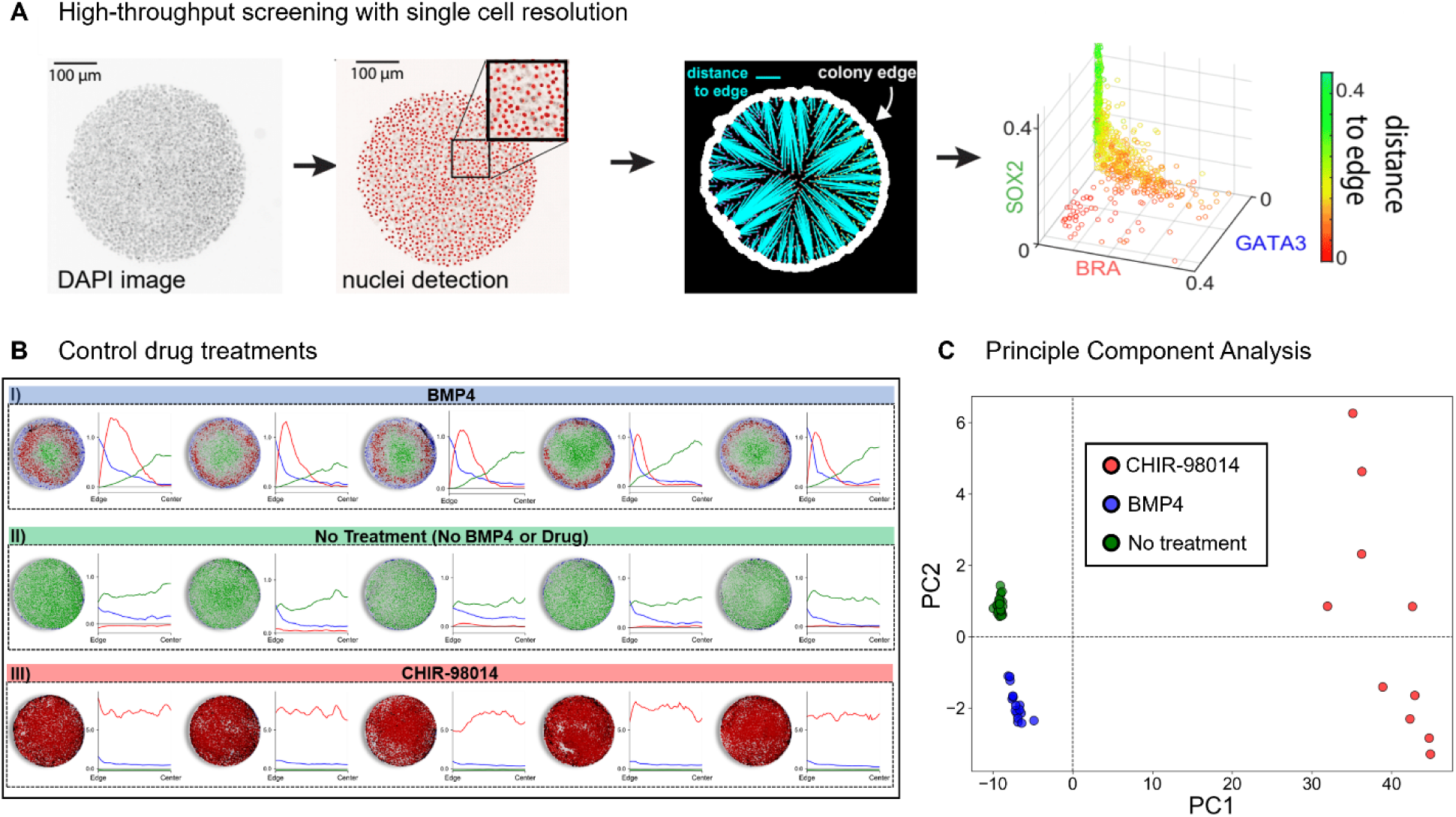
Automated image processing and quantification of 2D gastruloid patterning with single-cell resolution. (A) Schematic of the automated image processing pipeline showing single-cell resolution detection. DAPI-stained images of 2D gastruloids are processed to detect nuclei, allowing precise measurement of each cell’s location and expression levels within the colony, rather than averaging staining across radial bins. (B) Images and corresponding vector representations of control treatments. (I) BMP4 treatment leads to canonical radial patterning with GATA3 at the edge, BRA in the intermediate region, and SOX2 concentrated at the center. (II) No treatment (No BMP4 or drug) results in colonies maintaining pluripotency with high SOX2 expression throughout. (III) CHIR-98014 treatment induces BRA overexpression across the colony, disrupting canonical patterning. (C) Principal Component Analysis (PCA) of the 150-dimensional vector data demonstrating the ability to reliably separate control treatments (BMP4, No Treatment, CHIR-98014).

**Figure S2:**
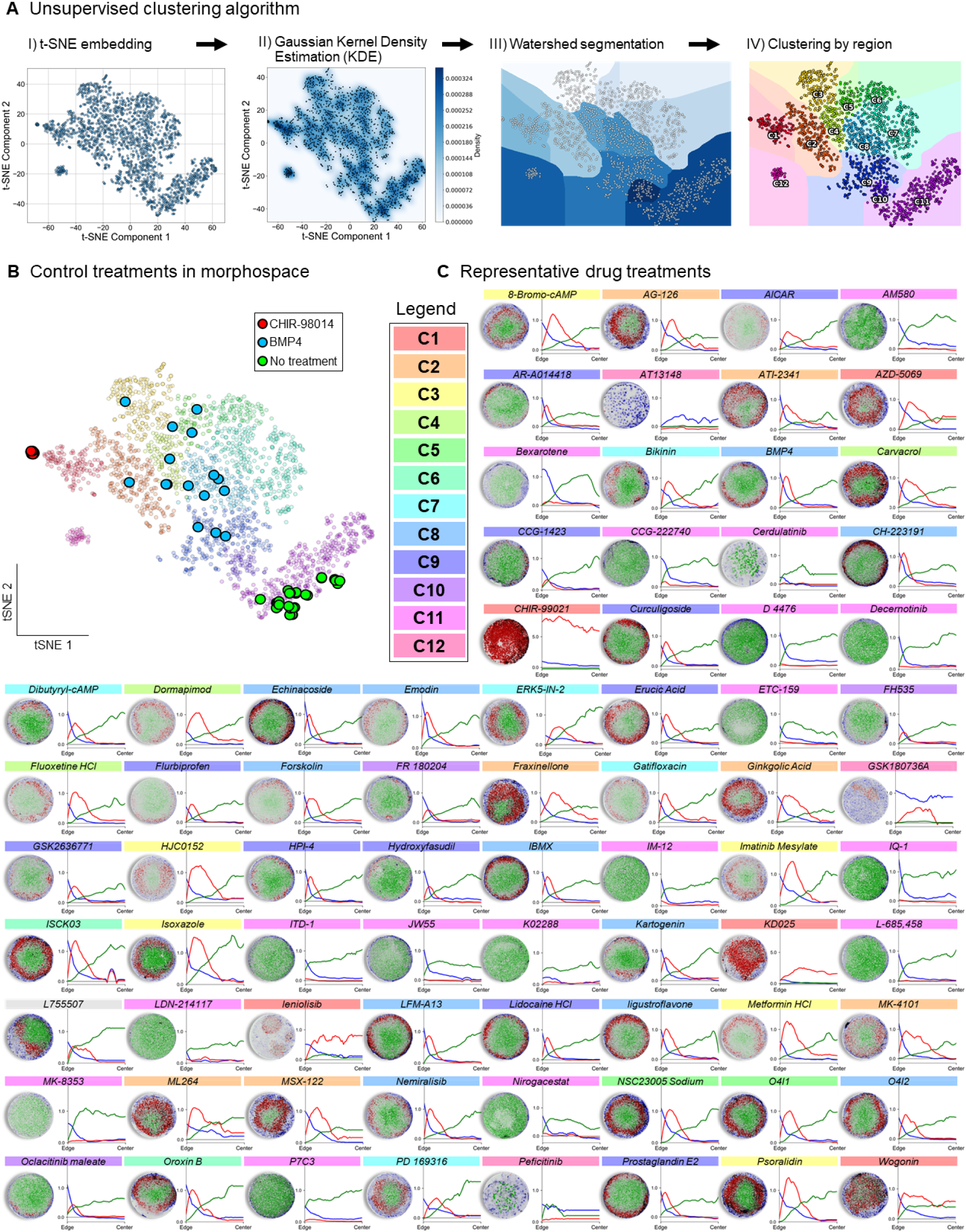
Unsupervised clustering algorithm and mapping of drug treatments in 2D gastruloid morphospace. (A) Schematic of the unsupervised clustering pipeline, which includes: (I) t-SNE embedding of high-dimensional data, (II) Gaussian Kernel Density Estimation (KDE) to smooth the data, (III) watershed segmentation applied to the smoothed density landscape, and (IV) clustering of colonies by region. (B) Location of control treatments (BMP4, CHIR-98014, No Treatment) within the 2D gastruloid morphospace, showing their distinct clustering patterns that validate the robustness of the clustering method. (C) Representative drug treatments mapped onto morphospace, with each drug’s corresponding cluster color-coded at the top. The displayed colonies and their radial intensity plots for GATA3, BRA, and SOX2 highlight the diverse morphological phenotypes induced by different compounds.

**Figure S3:**
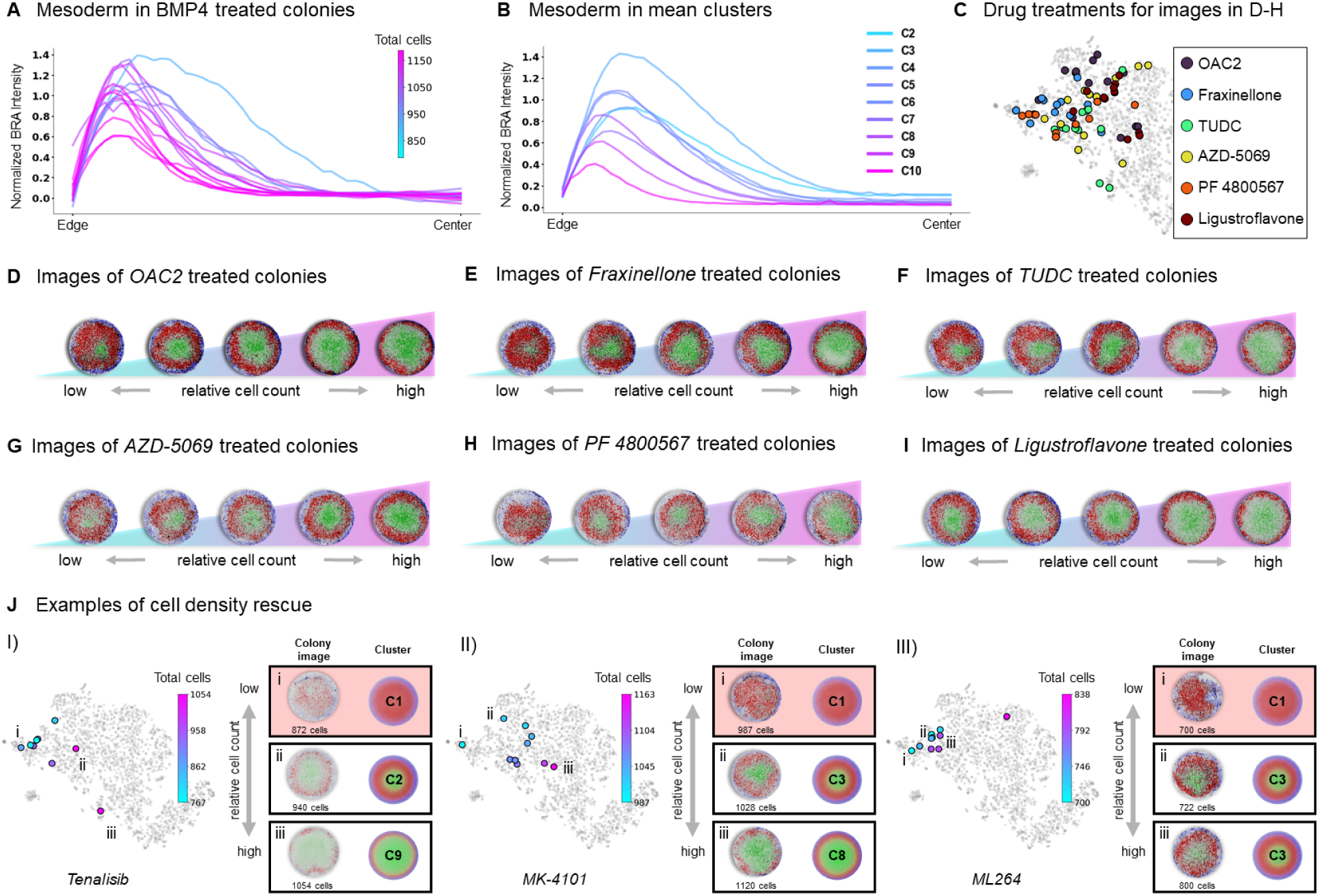
Impact of cell density on mesoderm patterning and rescue of failure modes in 2D gastruloids. (A-B) Mesoderm (BRA) staining intensity profiles across the colony in BMP4-treated controls (A) and mean clusters (B), colored by cell density. (C-I) Sample drug treatments demonstrating that the observed density-dependent patterning is not limited to BMP4 controls. (C) Location of colonies treated with OAC2, Fraxinellone, TUDC, AZD-5069, PF 4800567, and Ligustroflavone within the morphospace. (D-I) Images of treated colonies arranged by relative cell count, showing consistent trends of decreased mesoderm presence with increasing cell number, emphasizing that cell density dominates the patterning outcomes across different treatments. (J) Additional examples of cell density-based rescue and exacerbation of gastrulation failure modes. (I) High-density conditions can rescue colonies from failure mode clusters, as seen with Tenalisib-treated colonies transitioning from failure mode cluster C1 to canonical patterns as density increases. (II) MK-4101-treated colonies show that a low-density outlier causes the colony to drop into a failure mode. (III) Low-density conditions exacerbate failure modes, with ML264-treated colonies shifting into failure clusters as density decreases.

**Figure S4:**
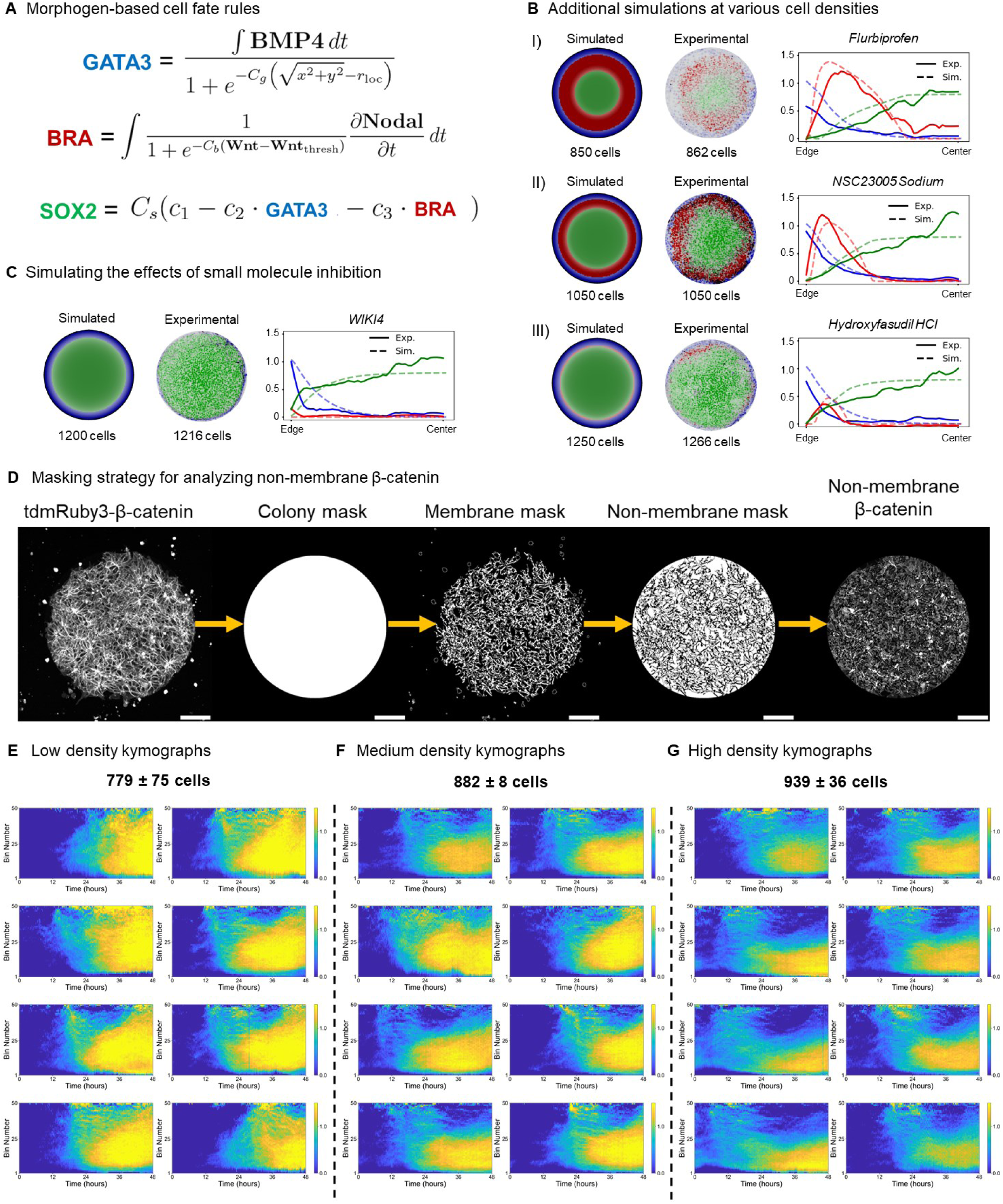
Morphogen-based cell fate determination and quantification of Wnt dynamics in 2D gastruloids. (A) Specific morphogen-based cell fate rules used in the simulations, detailing the equations governing the influence of BMP4, Wnt, and Nodal on GATA3, BRA, and SOX2 expression levels, as well as the role of SOX2 stability parameters. For more details, see Methods. (B) Additional simulations versus experimental comparisons at various cell densities (850, 1050, and 1250 cells), demonstrating the model’s ability to replicate patterns observed in the screen. Simulated (left) and experimental images (right) with corresponding line plots highlight the spatial distributions of cell fate markers (GATA3, BRA, SOX2). (C) Validation of the model’s ability to simulate the effects of small molecule inhibition. Shown is a colony treated with WIKI4, a Tankyrase inhibitor that blocks Wnt/β-catenin signaling. In the simulation, the cell-fate rules remained unchanged, but Wnt autoactivation was set to 0, accurately capturing the observed effects of WIKI4 on patterning. Scale bar: 100 µm. (D) Masking strategy for analyzing non-membrane β-catenin, showing the sequential extraction from tdmRuby3-β-catenin images to isolate active signaling components, separating non-membrane β-catenin from the membrane-bound pool. (E-G) Individual kymographs grouped by cell density ranges (low, medium, high) that were used to construct the mean kymographs shown in Figure 4F.

**Figure S5:**
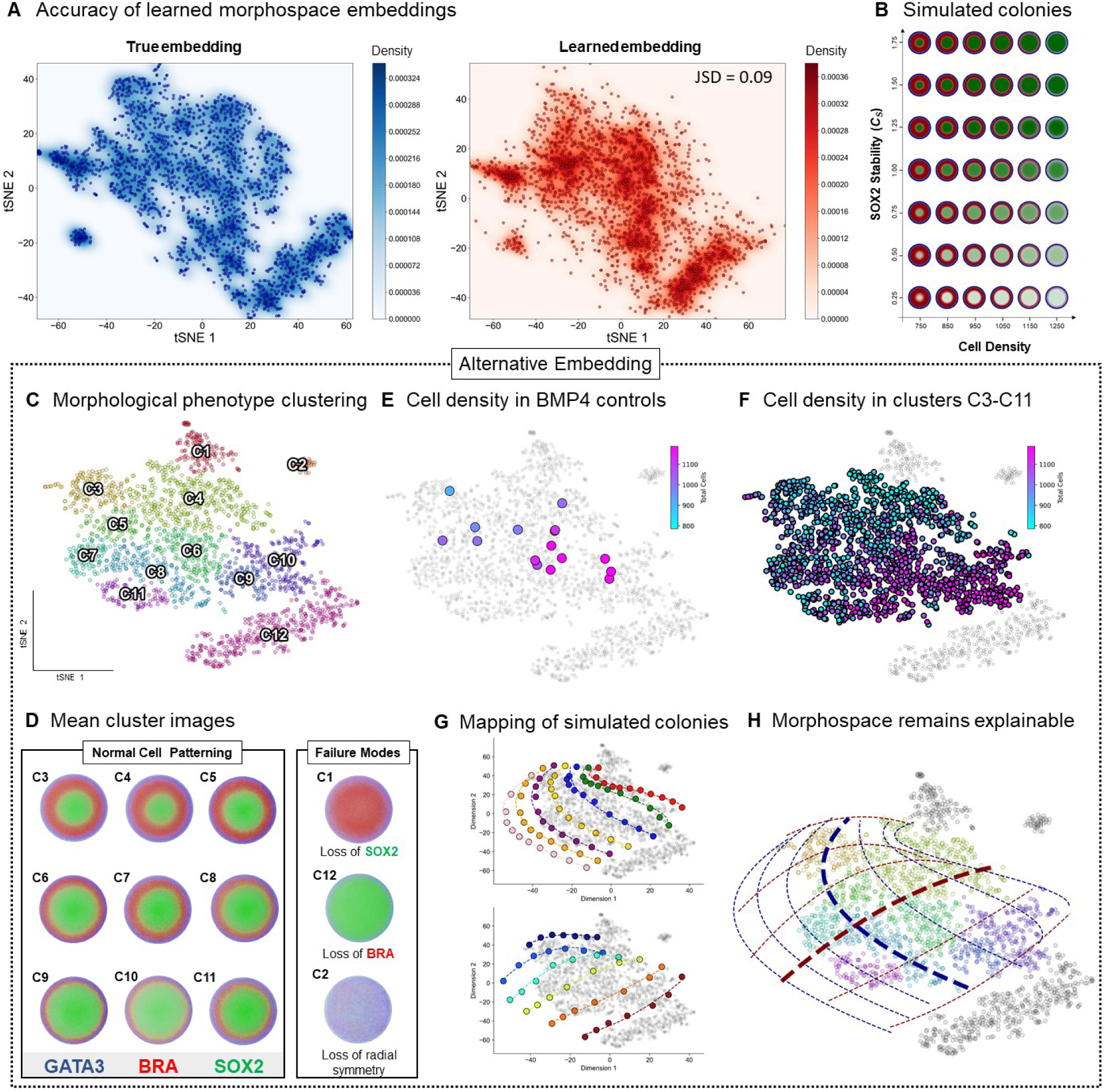
Validation of neural network accuracy in learning the low-dimensional embedding and consistency of morphospace explainability under alternative embedding conditions. (A) Density distributions of the true embedding (left, blue) derived from experimental data and the learned embedding (right, red) generated by the neural network. The low Jensen-Shannon Divergence (JSD = 0.09) between the distributions confirms that the learned embedding accurately replicates the experimental morphospace. (B) Simulated colonies showing changes in patterning as a function of cell density and SOX2 stability. (C-H) Construction of an alternative morphospace using a different seed for the t-SNE embedding, demonstrating robustness and consistency despite variations in hyperparameters. (C) The alternative morphospace retains the same number of clusters, though their spatial arrangement differs, confirming that clustering results are not specific to a single embedding. (D) Mean cluster images from the alternative embedding consistently capture canonical cell patterning and failure modes (e.g., loss of SOX2, loss of BRA, and loss of radial symmetry). (E-F) Impact of cell density on BMP4-treated controls and canonical patterning clusters (C3-C11) shows the same relationships as the original embedding. (G) Mapping of simulated colonies onto the alternative morphospace shows that cell density and SOX2 stability still define the two primary axes. (H) The morphospace remains explainable, with fitted polynomials still forming distinct axes, validating that observed trends and explainability are inherent to the data rather than artifacts of specific embedding parameters.

**Figure S6:**
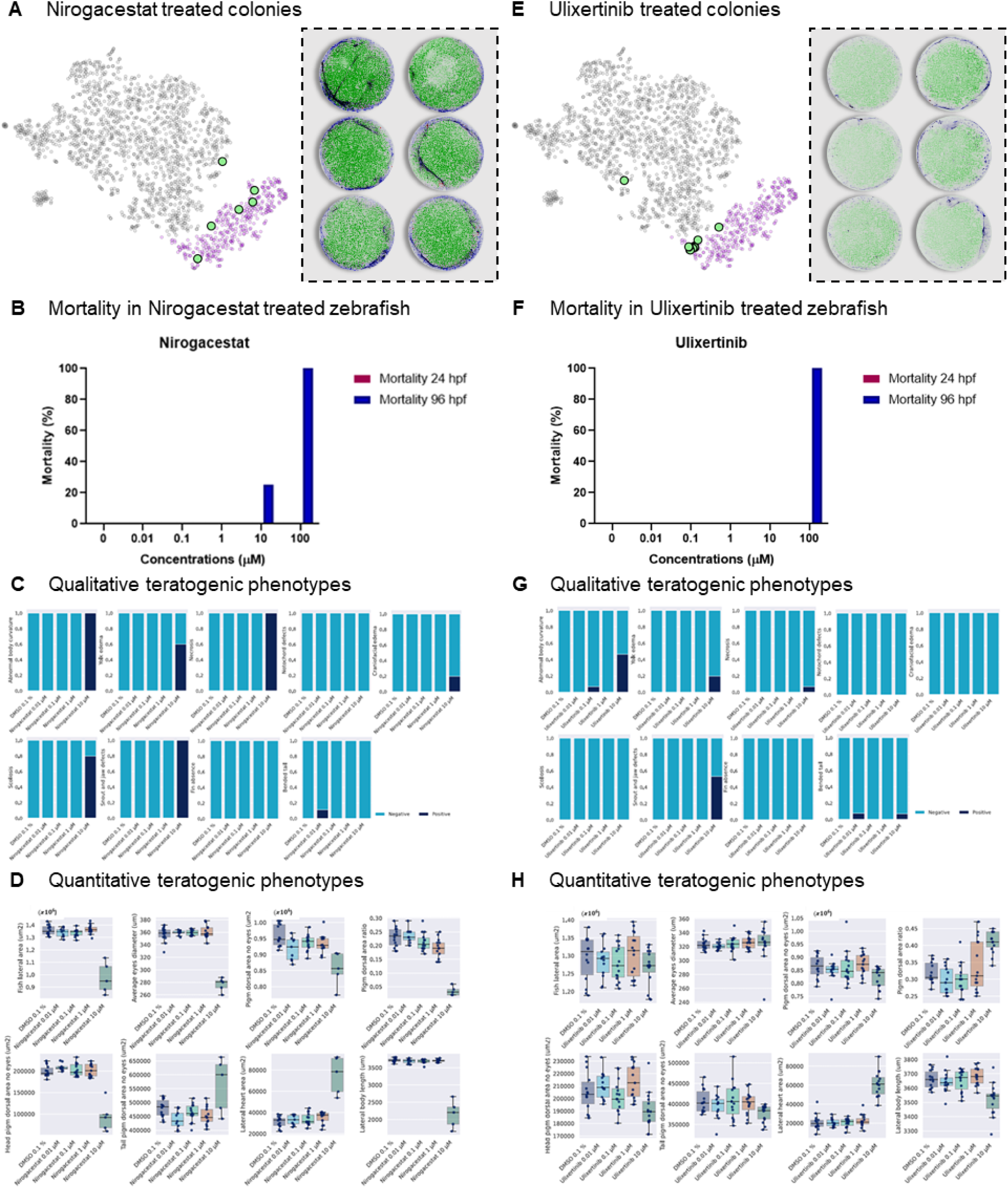
Teratogenic assessment of Nirogacestat and Ulixertinib in 2D gastruloids and zebrafish models. (A) Nirogacestat-treated colonies mapped in morphospace (left), highlighting their positioning within failure mode cluster C11 (purple), and representative images of the treated colonies (right), showing disrupted patterning consistent with potential teratogenic effects. (B) Mortality rates of zebrafish embryos treated with Nirogacestat at different concentrations, measured at 24 and 96 hours post-fertilization (hpf). (C) Qualitative teratogenic phenotypes observed in Nirogacestat-treated zebrafish, including abnormal body curvature, yolk edema, and other defects, with a high frequency of positive phenotypes at increased concentrations. (D) Quantitative teratogenic phenotypes in Nirogacestat-treated zebrafish, including lateral heart area and body length, showing significant alterations in comparison to controls, indicative of developmental toxicity. (E) Ulixertinib-treated colonies mapped in morphospace (left), demonstrating their clustering in C11, with representative images of the treated colonies (right). (F) Mortality rates of zebrafish embryos treated with Ulixertinib at varying concentrations, recorded at 24 and 96 hpf. (G) Qualitative teratogenic phenotypes observed in Ulixertinib-treated zebrafish. (H) Quantitative teratogenic phenotypes in Ulixertinib-treated zebrafish, showing significant deviations in metrics such as heart area and body length, further supporting the teratogenic potential of the compound.

## METHODS

### Cell Lines

All experiments were conducted using H9 (WA09) human embryonic stem cells obtained from WiCell (Cat No. WB66446). For the initial drug screen, H9 cells were utilized to assess the impact of various drug treatments on patterning outcomes. Additionally, a CRISPR-Cas9 tagged β-Catenin (β-Cat) line was developed using H9 cells specifically for the quantification of Wnt signaling dynamics, enabling the study of active, non-membrane-bound β-Catenin in live cells.

### Cell Culture

For routine maintenance, H9 cells were cultured on Corning Matrigel hESC-Qualified Matrix, LDEV-free (Corning, Cat #: 354277) coated dishes, and grown in mTeSR Plus feeder-free maintenance medium (STEMCELL Technologies, Cat #: 100-0276). Matrigel-coated dishes were prepared by coating them overnight at 4°C, followed by warming at room temperature for 1 hour before cell seeding.

## METHOD DETAILS

### Micropatterning

For micropattern cell culture, custom ordered 500 µm diameter micro-patterned 96 well glass bottom dishes (CYTOO Inc, Cat #: A500P650) were first coated with CellAdhere Laminin 521 (STEMCELL Technologies, Cat #: 200-0117) at 10 ug/mL for 2 hours at room temperature. Established protocols for 2D gastruloid patterning were used.^28^ Briefly, the wells are serially diluted with ice-cold calcium and magnesium free PBS (Thermo Fisher Scientific, Cat #: J67802.K2). Cells already resuspended in growth medium with 5 μM ROCK inhibitor (RI), Y-27632 (STEMCELL Technologies, Cat #: 72307) are then immediately plated upon removal of PBS. For cell seeding onto the micropattern, cells growing on Matrigel are lifted using ACCUTASE (STEMCELL Technologies, Cat #: 07920). Cells are centrifuged and 5x10^5^ cells per well are resuspended in RI containing media and added to the micropatterned plate. After 4 hours the medium is replaced with mTeSR plus without ROCK inhibitor and incubated overnight.

### Drug Screen

For the drug screen, we utilized the Stem Cell Signaling Compound Library (Selleck Chemicals, Cat #: L2100), containing 210 distinct drug conditions. Each compound was provided at an initial concentration of 10 mM, either dissolved in DMSO or water, based on solubility requirements. The compounds were stored and handled according to the manufacturer’s recommendations to maintain stability and activity. To prepare the working solutions, the compounds were aliquoted into 96-well plates and diluted to a final concentration of 10 μM in mTeSR Plus medium supplemented with BMP4 (STEMCELL Technologies, Cat #: 78211.1) at 50 ng/mL and without RI. Following the initial preparation of the micropatterned colonies, as described in the Micropatterning section, the cells were incubated overnight in mTeSR Plus medium. The next day, the media was carefully aspirated, and the BMP4 + drug mixture was added to the micropatterned wells, exposing the cells to both the differentiation cue (BMP4) and the specific drug conditions simultaneously. The plates were then incubated at 37°C with 5% CO₂ for 48 hours, allowing the cells to respond to the combined BMP4 and drug treatment.

### Immunofluorescence

Wells are fixed with 4% paraformaldehyde and rinsed twice with PBS then permeabilized with -80 °C ethanol for two minutes. Wells are then incubated with primary antibody at 4°C overnight (For antibodies see Key Resources Table), washed three times in PBS for 5 minutes each, and incubated with secondary antibody and DAPI nuclear counterstain (Thermo Fisher Scientific, Cat #: D3571) for 1 hour at room temperature before being washed 3 times with PBS.

### CRISPR-Cas9 fluorescent tagging

For CRISPR-Cas9 editing in H9 human embryonic stem cells, we utilized Lipofectamine Stem Transfection Reagent (Thermo Fisher Scientific, Cat #: STEM00015), to transfect a plasmid expressing a gRNA targeting the N-terminus of the *CTNNB1* gene, a homology template with ∼180bp of both upstream and downstream homology to the targeted insertion site flanking the tdmRuby2 construct, and the Cas9 enzyme (IDT) according to manufacturer’s directions. Edited cells were passaged into a single cell suspension and sorted, into ROCKi (5 µM in mTeSR), for tdmRuby2 positive clones using a Sony SH800 FACS system. Clones we isolated and expanded prior to verification of successful edition through PCR followed by Sanger sequencing.

### Confocal imaging of fixed 2D gastruloids from drug screen

All live and fixed cell imaging experiments were carried out using a Nikon W2 SoRa spinning-disk confocal microscope equipped with incubation chamber maintaining cells at 37 °C and 5% CO2. We imaged in the four channels corresponding to DAPI, Alexa488, Alexa555 and Alexa647 conjugated antibodies. Images were exported as .tiff files and analyzed using the custom MATLAB and Python software, enabling quantification of the spatial distribution and intensity of cell fate markers across various drug treatments.

### Live-cell confocal imaging

Cells were seeded at varying densities onto micropatterned plates using the CellVoyager CQ1 Benchtop High-Content Analysis System (Yokogawa Electric Corporation). Specifically, 5 × 10^5^ cells were resuspended in ROCK inhibitor-containing media and added to each well of the micropatterned plate. To achieve variation in initial seeding density, cells were diluted to 10%, 25%, 50%, 75%, and 100% concentrations (1:10, 1:4, 1:2, 1:1.33, and undiluted, respectively), with one well used for each condition. Eight colony positions were selected within each well using the CQ1 software, and these positions were saved in the protocol to allow for consistent imaging of the same locations after subsequent fixing and staining. The CQ1 system is equipped with an incubation chamber maintained at 37°C and 5% CO2 throughout the 48-hour experiment. Time-lapse imaging was performed with z-stacks acquired every 15 minutes using brightfield and 561 nm laser illumination. Laser autofocus was enabled to maintain image sharpness and consistency throughout the experiment. Images were automatically saved as .tiff files and exported for analysis. Post-acquisition, the images were processed and analyzed using custom Python software.

### Confocal imaging of fixed 2D gastruloids with varying density

Imaging was conducted using the same CellVoyager CQ1 Benchtop High-Content Analysis System to maintain consistency with the live-cell imaging setup. The previously saved positions from the live-cell imaging protocol were used to ensure that the same colonies were re-imaged, allowing for direct comparisons between live-cell dynamics and fixed-cell staining outcomes. Z-stacks were acquired for each of the four channels corresponding to DAPI, Alexa488, Alexa555, and Alexa647. All images were captured using the same incubation settings, with 37°C and 5% CO2 to prevent temperature-related artifacts during imaging. Laser autofocus was enabled for consistent focal plane alignment. Images were exported as .tiff files and analyzed using the custom MATLAB and Python software, enabling quantification of the spatial distribution and intensity of cell fate markers across varying seeding densities.

### Zebrafish Embryo Teratogenicity Assay and Phenotypic Analysis

All experiments were conducted using wild-type AB zebrafish (*Danio rerio*) maintained at 28 ± 1 °C with a 14-hour light and 10-hour dark cycle. Fertilized zebrafish embryos were collected in E3 1X medium (60X stock is prepared by dissolving 34.8 g NaCl, 1.6 g KCl, 5.8 g CaCl_2_·2H_2_O, 9.78 g MgCl_2_·6H_2_O in 2L H_2_O and adjusting the pH to 7.2 using NaOH) in Petri dishes, and abnormal or unfertilized embryos were discarded at 6 hours post-fertilization (hpf). Healthy embryos were grown up to 96 hpf and exposed to ulixertinib, nirogacestat, and isotretinoin at five concentrations (0.01 µM, 0.1 µM, 1 µM, 10 µM, 100 µM) starting at 6 hpf. 4-Diethylaminobenzaldehyde (DEAB) (MilliporeSigma, Cat #: D86256-100G) was used as a positive control at concentrations of 0.1 µM, 0.3 µM, 1 µM, 3 µM, and 10 µM.

Mortality was assessed at 24 h and 96 h, and larvae were imaged at 96 hpf using the automated VAST system (Union Biometrica). Dead, unhatched, and incorrectly detected larvae were excluded from analysis, resulting in a variable number of larvae analyzed per condition. Teratogenic effects were quantified by assessing both qualitative and quantitative phenotypes. Qualitative phenotypes included body curvature abnormalities, yolk edema, necrosis, and craniofacial defects, while quantitative phenotypes such as lateral area, eye diameter, pigmentation, and body length were measured using FIJI software. Continuous data were transformed to binary outcomes based on interquartile range thresholds.

LC50 and EC50 values, representing the concentration at which 50% of mortality is observed and the concentration at which 50% larvae population show a teratogenic phenotype, respectively, were calculated using dose-response models via Proast software, allowing for the determination of the teratogenic index (TI) as the ratio between LC50 and EC50 for the most sensitive phenotype. The Lowest Observed Effect Concentration (LOEC) was identified for each compound, with effects within 20% of biological range considered non-significant.

### Image Analysis

All image analysis was performed using custom software written in MATLAB and Python. The quantification of each imaged colony was performed in MATLAB using the same general workflow: background subtraction > dilation > nuclei detection > measurement. Subcellular segmentation of nuclear fluorescence was performed using DAPI brightness, size, and circularity to mask nuclei. Mean fluorescent intensity of regions of interest were measured and subsequently processed.

### Patterning quantification (MATLAB)

We developed in-house MATLAB software to extract intensity and positional intensity of imaged colonies. We first extract the DAPI image from the stack and rescale it into a binary image. Prior to true nuclei detection, we run the image through a background noise clean up algorithm, which is described as the following: using a morphological structuring element method, we define the nuclei criteria for detection. We then use a standard nuclei detection and enhancement pipeline, fill, dilation with our structuring element, followed by a final fill to get a grain free mask of the nuclear image channel. We lastly use MATLAB’s connected component detector (bwconncomp) to extract all nuclei positional and count information. With the generated mask, we go through a secondary segmentation step where once again use a morphological structuring element to enhance the binarized nuclei image followed by a regional maxima filter to extract the final nuclei positions. Intensity data is extracted by masking the generated segmentation across the imaged channels and finding the intensity values for each respective germ layer stain. To extract colony edge information, we can find the centroid information of all the nuclei before using a boundary detection algorithm to detect the edges of the colony along with each cell’s nearest border. Using the location data, we establish 50 bins that run from the center of the colony to the edge and fill each with the average intensity of cells in each bin for each channel. Lastly, we export both the raw intensity information and the binned intensity information.

### Dimensionality reduction (Python)

To extract trends from the imaged data set, we first apply robust scaling to the bin-wise data, normalizing each parameter within specific ranges (GATA, BRA, SOX) based on the 25th and 75th percentiles, and adjusting for each plate independently. This normalization is implemented using the pandas and numpy libraries, where numpy.percentile is used to calculate the 25th and 75th percentiles, and the data is scaled using scikit-learn’s RobustScaler. Normalizing by plate is crucial to account for plate-to-plate variability, ensuring that differences in experimental conditions or batch effects do not confound the analysis. This scaling method adjusts the data relative to the interquartile range (IQR), effectively minimizing the influence of outliers and centering the data on its median distribution. After scaling, we apply t-Distributed Stochastic Neighbor Embedding (t-SNE) using the TSNE function from the sklearn.manifold module to visualize high-dimensional data in a 2D space. The t-SNE output is refined using a custom unsupervised clustering algorithm that incorporates Kernel Density Estimation (KDE) with scipy.stats.gaussian_kde followed by watershed segmentation from skimage.segmentation, specifically using watershed and peak_local_max. This approach dynamically identifies distinct morphological phenotypes without requiring a predetermined number of clusters, distinguishing it from conventional clustering algorithms like k-means (sklearn.cluster.KMeans), Gaussian Mixture Models (sklearn.mixture.GaussianMixture), and spectral clustering (sklearn.cluster.SpectralClustering). The segmentation labels allow us to map and visualize clusters, enhancing the interpretability of the morphospace and identifying trends that characterize the different colony phenotypes observed in the experiments. Visualizations of the clusters and density surfaces were generated using matplotlib.pyplot and seaborn, and scatter plots were annotated with matplotlib.patheffects to enhance clarity.

### Quantifying Wnt signaling (Python)

To quantify active Wnt signaling, specifically the distribution of β-catenin within the cells, we developed a custom image processing algorithm to differentiate the active, non-membrane β-catenin from the inactive, membrane-bound pool. The algorithm was implemented using Python with the OpenCV and scikit-image libraries for image processing, along with numpy for numerical calculations. The processing pipeline began with reading the .tiff images using cv2.imread, followed by background subtraction to enhance the signal of the β-catenin staining. The core of the masking strategy involved segmenting the images to separate the membrane-bound and non-membrane pools of β-catenin. Initially, a Gaussian blur was applied using cv2.GaussianBlur to reduce noise. Then, adaptive thresholding (cv2.adaptiveThreshold) was used to generate a binary mask of the cell membrane, highlighting the high-intensity areas typically corresponding to membrane-bound β-catenin. For the non-membrane (active) β-catenin, the algorithm generated a complementary mask by dilating the initial binary mask with cv2.dilate and then subtracting this membrane mask from the original signal using cv2.subtract. This step effectively removed the membrane-bound signal, isolating the active β-catenin pool within the cytoplasm and nucleus. To further refine this mask, morphological operations such as cv2.erode were used to minimize any residual membrane signal bleed-through, ensuring that only cytoplasmic and nuclear β-catenin contributed to the final quantification. The extracted active β-catenin regions were then quantified by calculating the mean fluorescence intensity using numpy operations on the masked image arrays, providing a direct measurement of Wnt signaling activity. The results were visualized and statistically analyzed using matplotlib.pyplot and pandas.

### Generation of kymographs (Python)

To generate kymographs, a custom image processing pipeline was implemented to quantify the temporal dynamics of β-catenin (B-cat) signaling across multiple colony positions and time points. The process involved loading time-lapse images and applying a series of binning and quantification steps to create visual representations of spatial signaling changes over time. Images were first processed using the masking strategy previously described, where the active (non-membrane) β-catenin signal was isolated by subtracting the membrane-bound component. For each frame, this process was conducted using adaptive thresholding and morphological operations (cv2.threshold, cv2.dilate, and cv2.subtract) to create masks that accurately distinguished active signaling regions. Subsequently, the extracted non-membrane images were analyzed using a radial binning approach. Specifically, each colony was segmented into 50 concentric radial bins, defined by dividing the detected radius of the colony into equal segments using numpy.linspace. For each bin, mean fluorescence intensities were calculated by applying circular masks (cv2.circle) to the non-membrane β-catenin image, focusing on regions between each consecutive pair of radii. This quantification process was iterated over the entire set of frames, generating time-series data for each bin. The resulting kymographs were plotted using matplotlib.pyplot.imshow, providing a visual matrix where the x-axis represented time (in hours) and the y-axis represented the radial position of the bins. Color-coded intensity values indicated the level of active β-catenin signaling, allowing for clear visualization of temporal and spatial trends within each colony.

### Simulation of Reaction-Diffusion Models (Python)

The reaction-diffusion models were simulated using a custom Python script to capture the dynamic interactions of signaling pathways in the system. The simulation involved creating a 2D spatial grid to represent the environment and updating the concentrations of key signaling molecules over time. The script utilized Python libraries such as numpy for numerical computations, matplotlib for static and animated visualizations, and scipy for implementing diffusion and reaction processes. matplotlib.pyplot was used to generate heatmaps and plots of concentration profiles, while matplotlib.animation.FuncAnimation was employed to create time-lapse animations of the simulated morphogen distributions. Standard numerical techniques were applied to compute diffusion, reaction, and degradation processes, updating the system in small time steps. Custom functions handled boundary conditions and interactions between signaling molecules.

### Simulation of Cell Fate (Python)

Cell fate simulations were conducted using custom Python scripts designed to integrate the dynamics of morphogen signaling with rules for determining cell fate outcomes. The Python libraries used included numpy for numerical computations, matplotlib for visualizing data, and PIL (Python Imaging Library) for image manipulation and compositing. The process involved calculating fate maps based on our cell fate equations to determine the expression levels of GATA3, BRA, and SOX2 proteins. Visualization of fate maps was performed by generating heatmaps of normalized expression levels using custom color maps to represent each fate (GATA, BRA, and SOX). Separate images were created for each morphogen, and a composite image was generated to illustrate the combined spatial distribution of all fates within the colony. Additionally, radial intensity profiles were computed by binning the colony into concentric rings and averaging the protein expression levels within each ring.

### Neural Network-Based Approximation of t-SNE Embedding (Python)

To capture trends in high-dimensional data and map them into a low-dimensional space, we implemented a neural network-based approximation of the t-SNE embedding function. While there is no direct inverse solution to t-SNE, we constructed an approximate function, 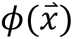, to predict the low-dimensional embedding of high-dimensional data points. This approximation was achieved by training a 4-layer neural network with ReLU activation functions to minimize the difference between the predicted and actual locations in the low-dimensional space generated by the original t-SNE. High-dimensional data vectors describing the bin-wise distribution of cell fate markers were used alongside their corresponding t-SNE embeddings.

Data preprocessing involved loading the datasets, filtering for numeric columns, and converting them to PyTorch tensors. We split the data into training and validation sets using an 80/20 split and created TensorDatasets and DataLoaders for efficient batch processing during training. The neural network consisted of three fully connected layers with decreasing dimensions (128, 64, and 2 neurons, respectively), reflecting the desired 2D embedding output.

Training was conducted over 10,000 epochs using the Adam optimizer with a learning rate of 0.001, coupled with mean squared error loss to quantify the difference between predicted and true low-dimensional embeddings. A learning rate scheduler with a ReduceLROnPlateau strategy was employed to adapt the learning rate based on validation loss improvements. Early stopping was implemented with a patience of 50 epochs to prevent overfitting, saving the best-performing model during training.

To evaluate the accuracy of 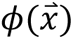, after training, the model was used to predict low-dimensional embeddings from the original high-dimensional data. Predictions were visualized alongside the original t-SNE embeddings, with density surfaces and scatter plots generated using Matplotlib and Seaborn. The primary quantitative measure employed to compare the predicted and actual embeddings was Jensen-Shannon Divergence (JSD), which assessed the similarity between the predicted and true density distributions in the low-dimensional space.

To further interpret the structure of the predicted low-dimensional embeddings, polynomial fitting was employed to visualize the “axes” of variation across different experimental conditions. Specifically, we projected simulated colonies, which were generated across multiple cell densities (ranging from 750 to 1250 cells) and varying stability values (C_s_ values ranging from 0.25 to 1.75), into the low-dimensional space using the neural network model. By predicting the low-dimensional embeddings for these simulated colonies, we identified trends within these embeddings by fitting polynomials to the observed trajectories.

Polynomial fitting was conducted using the numpy.polynomial.Polynomial class, enabling us to approximate the trajectory of data points in the reduced-dimensional space. Different polynomial degrees were applied depending on the variability and complexity of the data clusters: higher-order polynomials (up to third degree) were used to capture non-linear trends in groups with higher variability, while lower-order polynomials were applied where simpler, more linear relationships were evident. This fitting process facilitated the visualization of how specific experimental parameters, such as cell density and stability values, influenced the positioning of data points within the low-dimensional space, effectively revealing the “axes” along which these factors varied.

## Mathematical Modeling

### Reaction-diffusion model for morphogen spread

Inspired by recent studies examining BMP-Wnt-Nodal dynamics in 2D gastruloids,^41^ we developed a simplified reaction-diffusion model tailored to our experimental observations. The model specifically addresses the influence of cell density on mesoderm patterning, as we observed that lower cell density shifts the mesoderm peak inward and causes the mesoderm region to expand outward. This density-dependent modulation was incorporated into our model, accounting for the dynamic interactions between BMP, Wnt, and Nodal, and aligning with observed spatial and temporal morphogen patterns.

BMP was modeled with the following equation:

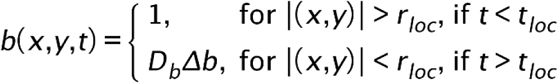

The BMP signaling dynamics are modeled as a diffusive species within a localized region, *r_loc_*, which varies based on cell density. Initially, BMP concentration is set to 1 across the entire domain, representing the uniform activation phase. After t_loc_, BMP diffusion is restricted within rloc, simulating the feedback inhibition observed experimentally, where BMP signaling diminishes centrally but remains active at the periphery.^34^ In this context, D_b_, represents the diffusion coefficient of BMP, r_loc_, denotes the radius defining the localized BMP diffusion modulated by cell density, and t_loc_ indicates the time point when BMP signaling becomes localized. The boundary for BMP signaling, r_loc_, is defined by *r_rad_* ∗ (1 − *e*^−*a*∗*d*^), where and r_rad_ is the radius of the stem cell colony. Here, *a* is a constant that scales the effect of cell density, and *d* represents the cell density; this formulation reflects our observation that the outermost cells spread more at lower densities, increasing the localized signaling boundary. The spread of Wnt was modeled by:

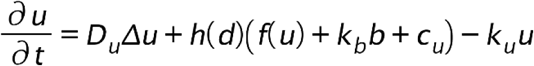

Where:

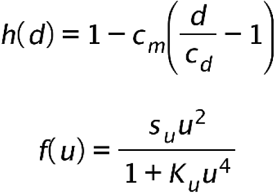

The Wnt signaling dynamics incorporate diffusion and autocatalytic activation, modulated by the density-dependent scaling function h(d). The term f(u) captures the autocatalytic production of Wnt, with inhibition effects included through a saturation term in the denominator to limit excessive Wnt production at high concentrations. Here, D_u_, is the diffusion coefficient of Wnt, h(d) is the density-dependent scaling function, s_u_ denotes the rate of Wnt autocatalytic production, K_u_ is the saturation constant controlling Wnt production, k_b_ represents the contribution of BMP to Wnt production, c_u_ is a constant source term for Wnt, and k_u_ is the degredation rate on Wnt. The scaling function h(d) modulates Wnt reaction terms based on cell density, where c_m_ is a scaling factor, _d_ is the cell density, and c_d_ is a constant representing the mean cell density from the screen. The spread of Nodal was modeled by:

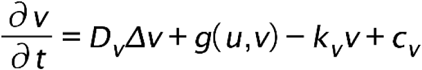

Where:

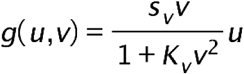

The Nodal signaling dynamics incorporate diffusion and production influenced by Wnt activity, as captured by the term g(u,v), which includes a saturation effect to reduce Nodal production at high concentrations. Unlike previous models, we chose not to model Nodal as autocatalytic due to evidence indicating that Nodal spreads primarily through a relay mechanism rather than an autocatalytic loop.^72^ In this context, D_v_, denotes the diffusion coefficient of Nodal, s_v_, is the rate of Nodal production influenced by Wnt, K_v_ is the saturation constant reducing Nodal production at high concentrations, c_v_ represents the constant source term for Nodal, and k_v_ is the degradation rate of Nodal. All parameter values used in the model can be found in Table S3.

### Morphogen-based cell fate determination model

We developed cell-fate determination rules based on the integration of BMP, Wnt, and Nodal signaling dynamics. Each fate map was generated by simulating the morphogen concentrations over time, followed by applying specific computational rules to map the spatial location of fate-markers, GATA3, BRA, and SOX2, throughout the colony. Specifically, GATA3 expression was modeled by:

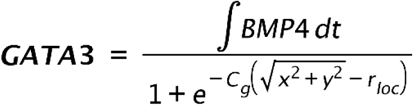

The expression of GATA3 is modeled as a response to the integral of BMP signaling over time, reflecting the cumulative impact of BMP exposure on cells.^37^ The denominator accounts for the preferential availability of transforming growth factor β receptors at the edge of the colony,^34^ with the steepness parameter modulating the influence of distance from the colony edge. Here, 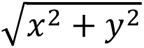 represents the radial position within the colony, r_loc_ defines the localized region where BMP signaling is most concentrated, and C_g_ adjusts the drop-off rate of GATA3 expression as distance increases from the colony edge increased. BRA expression was modeled by:

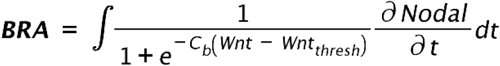

BRA expression is governed by the combined influence of Wnt and the rate of change of Nodal signaling, capturing the dynamics where Nodal alone does not induce mesoderm formation.^73^ Here, the Wnt concentration must reach a certain threshold, Wnt_thresh_ in order for the cells to respond to changes in the rate of Nodal concentration. Specifically, unlike BMP, which is concentration-dependent, our experimental observations indicate that many Nodal targets depend on the rate of concentration change rather than its absolute levels. Here, C_b_ represents the steepness of the BRA activation response to Wnt, Wnt_thresh_ denotes the threshold level of Wnt required for BRA expression, and 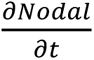 is the rate of change of Nodal concentration over time. SOX2 expression was modeled by:

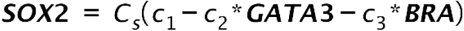

SOX2 expression is modeled as inversely proportional to the levels of GATA3 and BRA, aligning with our observations that SOX2 expression typically decreases as cells begin to differentiate. The stability constant, C_s_, modulates the persistence of SOX2 under varying conditions, reflecting our experimental findings that certain drug treatments allow cells to maintain high SOX2 expression despite concurrent BRA expression. In this equation, C_1_, C_2_, and C_3_ are constants that adjust the relative influence of GATA3 and BRA on SOX2 suppression. All parameter values used in the model can be found in Table S4.

## ADDITIONAL DETAILS

Our map of 2D gastruloid morphopsace results can be explored via the interactive website: https://max-wilson.mcdb.ucsb.edu/research/gastruloid-morphospace

